# DIRAS3 broadly suppresses signaling from diverse oncogenic KRAS hotspot variants through an inhibitory interface

**DOI:** 10.64898/2026.10.21.760748

**Authors:** Gamze Bildik, Hailing Yang, Weiqun Mao, Ritchel B. Gannaban, Viyaj K. Bhardwaj, Md Nasir Uddin, Sung Y. Jung, Mariaelena Pierobon, Alison Gomeiz, Alemayehu A. Gorfe, John F. Hancock, Christopher B. Marshall, Frank McCormick, Steven W. Millward, Junchen Liu, Robert C. Bast, Zhen Lu

## Abstract

Despite major advances in KRAS-directed therapies, broadly suppressing oncogenic KRAS signaling across diverse hotspot variants remains an unmet challenge. Here, we identify an endogenous inhibitory mechanism that suppresses signaling across the major oncogenic KRAS hotspot classes while largely sparing basal wild-type KRAS, revealing activation-state-selective inhibition. We demonstrate that the imprinted tumor suppressor DIRAS3 directly engages activated KRAS through a functionally defined inhibitory interface that can be molecularly mimicked to broadly inhibit KRAS-driven signaling. Using genetically controlled RASless reconstitution models, complementary cancer cell validation, structure-function analyses, molecular dynamics-based membrane topology modeling, and ultrastructural imaging, we show that dual lipid modification constrains DIRAS3 membrane topology, enabling an α5-encoded inhibitory interface to engage activated KRAS, disrupt signaling-competent membrane assemblies, and suppress downstream signaling. Structure-function analyses establish this interface as a critical determinant of DIRAS3-mediated KRAS inhibition. A conformationally constrained α5 mimetic recapitulates full-length DIRAS3 activity, broadly suppressing MAPK and PI3K signaling across multiple oncogenic KRAS variants, including Q61 mutants with reduced sensitivity to current tri-complex RAS(ON) inhibitors. Mechanistically, immunoprecipitation shows that the α5 mimetic reduces KRAS association with both RAF1 and p110α similarly across G12 and Q61 mutants, whereas RMC-7977 reduces inhibition of p110α association with Q61 KRAS. The α5 mimetic also disrupts KRAS nanocluster organization and inhibits tumor growth in vivo. These findings identify a targetable inhibitory interface within DIRAS3 and establish mimicry of an endogenous KRAS suppressor as a mechanistically distinct strategy for broad KRAS inhibition.

**Significance:** These findings establish that DIRAS3-mediated endogenous regulation extends across diverse oncogenic KRAS variants, including Q61X mutants, revealing an unrecognized principle of pan-KRAS regulation that can be recapitulated by a DIRAS3-derived peptide.

## Introduction

Activating mutations in KRAS are among the most prevalent oncogenic drivers in human cancer and promote sustained signaling through the MAPK and PI3K pathways^1–3^. Despite decades of efforts, direct pharmacologic inhibition of KRAS has proven challenging, due to its high affinity for GTP/GDP and the limited accessibility of ligandable pockets. Recent advances have yielded allele-specific covalent inhibitors targeting KRAS G12C and a new generation of RAS-ON inhibitors designed to engage the GTP-bound state of KRAS. Emerging evidence further suggests that active-state RAS inhibitors suppress signaling not solely through nucleotide-state recognition, but also through formation of inhibitory complexes that perturb effector engagement and destabilize signaling-competent KRAS assemblies^4^. While these approaches represent major therapeutic breakthroughs, their activity remains constrained by mutation specificity, adaptive resistance, and incomplete suppression of downstream signaling, particularly in tumors driven by Q61 substitutions.

Beyond nucleotide state and catalytic cycling, KRAS function is critically dependent on its spatial organization at the plasma membrane^5, 6^. Active, GTP-loaded KRAS assembles into dynamic nanoclusters that serve as signaling platforms for effector recruitment, RAF1 dimerization, and downstream signaling^7^. Recent studies further support the concept that signaling-competent KRAS exists within higher-order, effector-coupled membrane assemblies whose stability contributes to sustained pathway output^4, 8^. Disruption of KRAS membrane organization has therefore emerged as a potential strategy to block oncogenic signaling without direct targeting of the KRAS nucleotide-binding pocket.

DIRAS3 is an endogenous imprinted tumor suppressor of RAS^9–12^. As an atypical member of the RAS superfamily, DIRAS3 lacks intrinsic oncogenic activity yet preserves the conserved GTPase core architecture of canonical RAS proteins. Unlike classical RAS isoforms, DIRAS3 contains a unique 34-amino acid N-terminal extension that enables suppression of RAS function. DIRAS3 has been shown to bind directly to RAS and to disrupt RAS nanoclusters, but the mechanism by which it inhibits signaling-competent mutant KRAS is not completely understood.

Here, we identify a topology-dependent inhibitory interface within the endogenous tumor suppressor DIRAS3 that enables broad suppression of oncogenic KRAS signaling. Using RASless reconstitution systems, mutational dissection, molecular dynamics-based membrane topology modeling, and ultrastructural imaging, we demonstrate that dual lipid modification positions the DIRAS3 α5 region to engage active KRAS, disrupt signaling-competent nanoclusters, and attenuate downstream signaling. This mechanism preferentially suppresses oncogenic, but not wild-type, KRAS and is preserved across multiple oncogenic KRAS variants. Furthermore, we show that a minimal DIRAS3 α5-derived mimetic peptide is sufficient to recapitulate DIRAS3-mediated inhibition by disrupting KRAS association with both RAF1 and p110α across G12X and Q61X variants. These findings establish endogenous suppressor mimicry as a mechanistically distinct strategy for broad inhibition of oncogenic KRAS. p110α across G12X and Q61X variants. These findings establish endogenous suppressor mimicry as a mechanistically distinct strategy for broad inhibition of oncogenic KRAS.

## Results

### Membrane topology enables DIRAS3-mediated suppression of oncogenic KRAS

A mechanistic understanding of how DIRAS3 suppresses oncogenic KRAS requires a precise definition of the DIRAS3 structure that mediates KRAS engagement, regulates KRAS spatial organization at the plasma membrane, and couples nanocluster disruption to downstream signaling inhibition^8, 13–15^. We previously demonstrated that effective suppression of KRAS signaling by DIRAS3 requires both its N-terminal extension (NTE), which contains a myristoylation site, and its C-terminal CAAX motif for prenylation, implicating membrane association as a prerequisite for function^10, 11^. However, the structural features that govern KRAS engagement, spatial organization at the plasma membrane, and coupling to downstream signaling inhibition have remained incompletely defined.

To define the molecular basis for broad suppression of oncogenic KRAS by DIRAS3, we first sought to identify the structural determinants required for KRAS engagement and signaling inhibition. We generated a panel of DIRAS3 mutants, targeting N- and C-terminal regions, lipid modification sites, and predicted interaction surfaces **(Figure 1A-B and Supplementary Figure 1A)**. These include deletion of the N-terminal extension (ΔNTE; residues 35-229), deletion of the C-terminal region containing the CAAX motif (ΔCTE; residues 1-206), combined terminal deletions (ΔNCTE; residues 35-206), and point mutations disrupting N-terminal myristoylation (G2A), C-terminal prenylation (C226S), and targeted internal deletions and substitutions. All constructs were expressed in RASless murine embryonic fibroblasts (MEFs) rescued with oncogenic KRAS G12D, enabling direct evaluation of KRAS-dependent signaling in a uniform genetic background **(Figure 1C)**.

**Figure 1.**
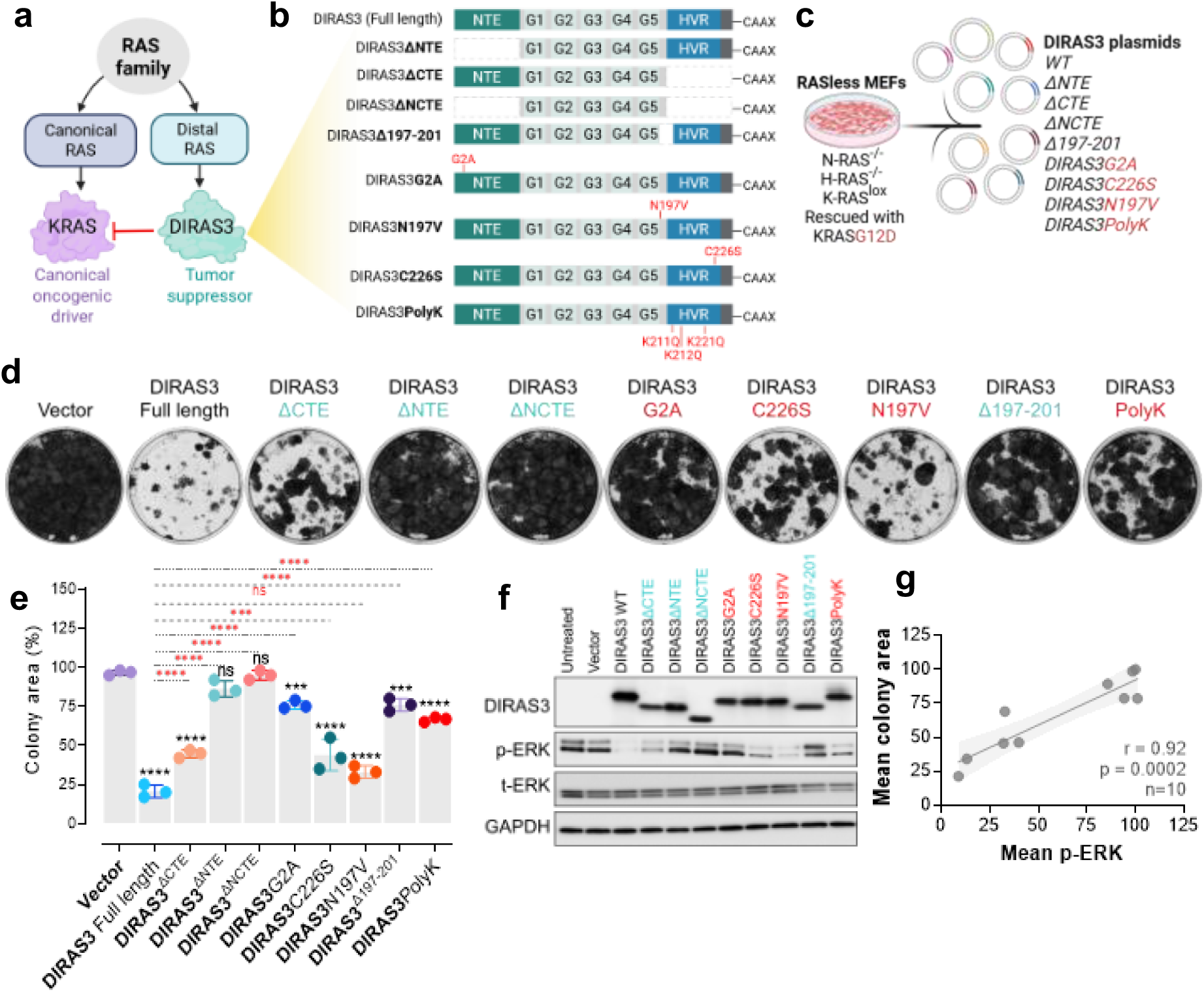
Structural dissection of DIRAS3 identifies determinants required for KRAS inhibition. **(a)** Schematic illustrating DIRAS3 as a unique tumor suppressor within the RAS superfamily and the overall study rationale. Structural dissection of DIRAS3 was performed to identify the inhibitory interface responsible for KRAS suppression. **(b)** Schematic of DIRAS3 deletion and point mutants targeting the N-terminal extension (ΔNTE), C-terminal extension (ΔCTE), combined termini (ΔNCTE), α5 helical region (Δ197-201 and N197V), and lipid modification sites (G2A and C226S), and the polybasic membrane-binding region (PolyK; K211Q/K212Q/K221Q). **(c)** Overview of the isogenic RASless MEF system rescued with KRAS^G12D^ and transfected with different DIRAS3 variants. **(d)** Representative images of colony formation assays following transient expression of WT or mutant DIRAS3 in KRAS^G12D^ RASless MEFs. Colonies were stained with Coomassie Brilliant Blue 10-14 days post-seeding. **(e)** Quantification of total colony area per well was measured using ImageJ and analyzed in GraphPad. Data are presented as mean ± SD from three independent experiments (n=3). Statistical significance was determined using two-way ANOVA. Comparisons to empty vector control (black) and WT DIRAS3 (red) are indicated. ns, p > 0.05; ***p < 0.001; ****p < 0.0001. **(f)** Immunoblots showing phospho-ERK1/2 levels following expression of WT or mutant DIRAS3 in RASless MEFs rescued with KRAS^G12D^. **(g)** Correlation analysis of clonogenic growth and ERK signaling activity. (Linear regression analysis, r = 0.92, p = 0.0002). Schematic in panels (a-c) created in BioRender. [https://BioRender.com/wwipwx3, https://BioRender.com/9geb100]

Wild-type DIRAS3 significantly impaired clonogenic growth **(Figure 1D-E)** and robustly suppressed ERK1/2 phosphorylation **(Figure 1F)**. Combined deletion of both termini fully abolished DIRAS3 function. Deletion of the N-terminal extension (ΔNTE) or elimination of the N-terminal myristoylation site (G2A) completely abrogated DIRAS3-mediated suppression of MAPK signaling and clonogenicity **(Figure 1D-F, Supplementary Figure 1B)**. In contrast, deletion of the C-terminal CAAX-containing region (ΔCTE) or mutation of the prenylation site (C226S) resulted in partial, but not complete, loss of inhibitory activity **(Figure 1D-F, Supplementary Figure 1B)**. Importantly, both ΔNTE and ΔCTE mutants retained plasma membrane association **(Supplementary Figure 1C)**, indicating that loss of function is not due to impaired membrane localization but instead may reflect disruption of the membrane topology required for DIRAS3 activity. These findings demonstrate that dual lipid anchoring is required for DIRAS3 function, likely by establishing the correct membrane orientation, with N-terminal myristoylation contributing disproportionately to effective KRAS suppression. This asymmetric dependence distinguishes DIRAS3 from canonical RAS family members, which rely predominantly on C-terminal CAAX-mediated prenylation for membrane association^16–18^. In addition, across the panel of DIRAS3 mutants, ERK inhibition strongly correlated with inhibition of clonogenic growth (r = 0.92), demonstrating that proliferative output is tightly coupled to MAPK activity in this system **(Figure 1G)**.

To confirm that these lipid modifications occur in cancer cells, we examined endogenous DIRAS3 modification in KRAS-driven tumor cell lines. Mass spectrometric analysis of DIRAS3 isolated from KRAS-mutant pancreatic ductal adenocarcinoma and low-grade serous ovarian carcinoma cell lines consistently identified N-terminal myristoylation at glycine 2 (with removal of Met1) and C-terminal prenylation at the CAAX motif **(Supplementary Figure 2A-C)**. These findings confirm that DIRAS3 undergoes dual lipid modification in cancer cells, providing a structural basis for stable plasma membrane association in oncogenic KRAS contexts and validating the biological relevance of the lipidation-dependent functional requirements observed in the mutant analyses.

We next examined whether DIRAS3 similarly suppresses KRAS-MAPK signaling in vivo using a pancreatic cancer xenograft model (AsPC-1) with Tet-inducible DIRAS3 expression. Following 8 days of doxycycline treatment, control tumors without DIRAS3 induction exhibited strong p-ERK staining, consistent with active oncogenic KRAS signaling. In contrast, tumor cells expressing DIRAS3 showed markedly reduced p-ERK levels, indicating suppression of MAPK pathway activity in vivo **(Supplementary Figure 2D)** and supporting the physiological relevance of DIRAS3-mediated suppression.

Together, these findings establish that full-length DIRAS3 requires dual lipid modification to achieve a membrane topology necessary for effective suppression of oncogenic KRAS signaling in vitro and in vivo.

### DIRAS3 encodes an α5 inhibitory interface that selectively destabilizes signaling-competent KRAS nanoclusters

Beyond membrane anchoring, we observed that deletion of the DIRAS3 α5-associated segment encompassing residues 197-201 (Δ197-201) profoundly impaired DIRAS3-mediated inhibition of ERK phosphorylation and clonogenic growth **(Figure 1D-F, Supplementary Figure 1B)**. This region lies within a predicted KRAS-interacting surface that is spatially aligned with the KRAS α5 helix^19^, suggesting a role in mediating KRAS engagement or perturbing KRAS organization. Although sequence similarity to KRAS is limited **(Supplementary Figure 1A)** and AlphaFold predictions do not strongly support a stable helix in isolation^20^, these analyses do not preclude a structured conformation in the context of the membrane or upon KRAS engagement. We interrogated residue N197 because it occupies a position analogous to D154 in KRAS, a critical determinant of the α4-α5 dimer interface implicated in KRAS signaling output^21^. However, substitution of the conserved N197 residue with valine (N197V) preserved most inhibitory activity, indicating that the loss of function following Δ197-201 deletion reflects disruption of local α5 structural integrity rather than mutation of a single side chain. Mutation of the poly-lysine cluster (K211Q/K212Q/K221Q) resulted in partial loss of inhibitory activity, consistent with a modulatory role for electrostatic membrane interactions.

To define the membrane topology underlying α5-dependent inhibition of KRAS, we performed all-atom molecular dynamics (MD) simulations of full-length, lipidated DIRAS3 on a model plasma membrane. The simulations showed a rapid decrease in the distance between the center-of-mass of the G-domain and lower leaflet phosphorous (P) atoms dropped from 45 Å to approximately ∼20 Å within the first microsecond, followed by stabilization at ∼15 Å **(Supplementary Figure 3A)**. Consistently, the root mean square deviation (RMSD) analysis of the G-domain Cα atoms indicated overall structural stability with limited conformational deviation over the simulation time course **(Supplementary Figure 3B)**. DIRAS3 rapidly adopted a stable membrane-bound conformation through dual N-terminal myristoylation and C-terminal prenylation **(Figure 2A)**. This topology positions the canonical effector-binding interface encompassing switch I and switch II oriented toward and partially occluded by the membrane, whereas the α5 helix projects away from the bilayer into the cytosol and remains solvent-exposed. These structural features further support a membrane topology that is unfavorable for canonical effector engagement through switch-region interactions while positioning the α5 region for potential protein-protein interactions at the membrane interface.

**Figure 2.**
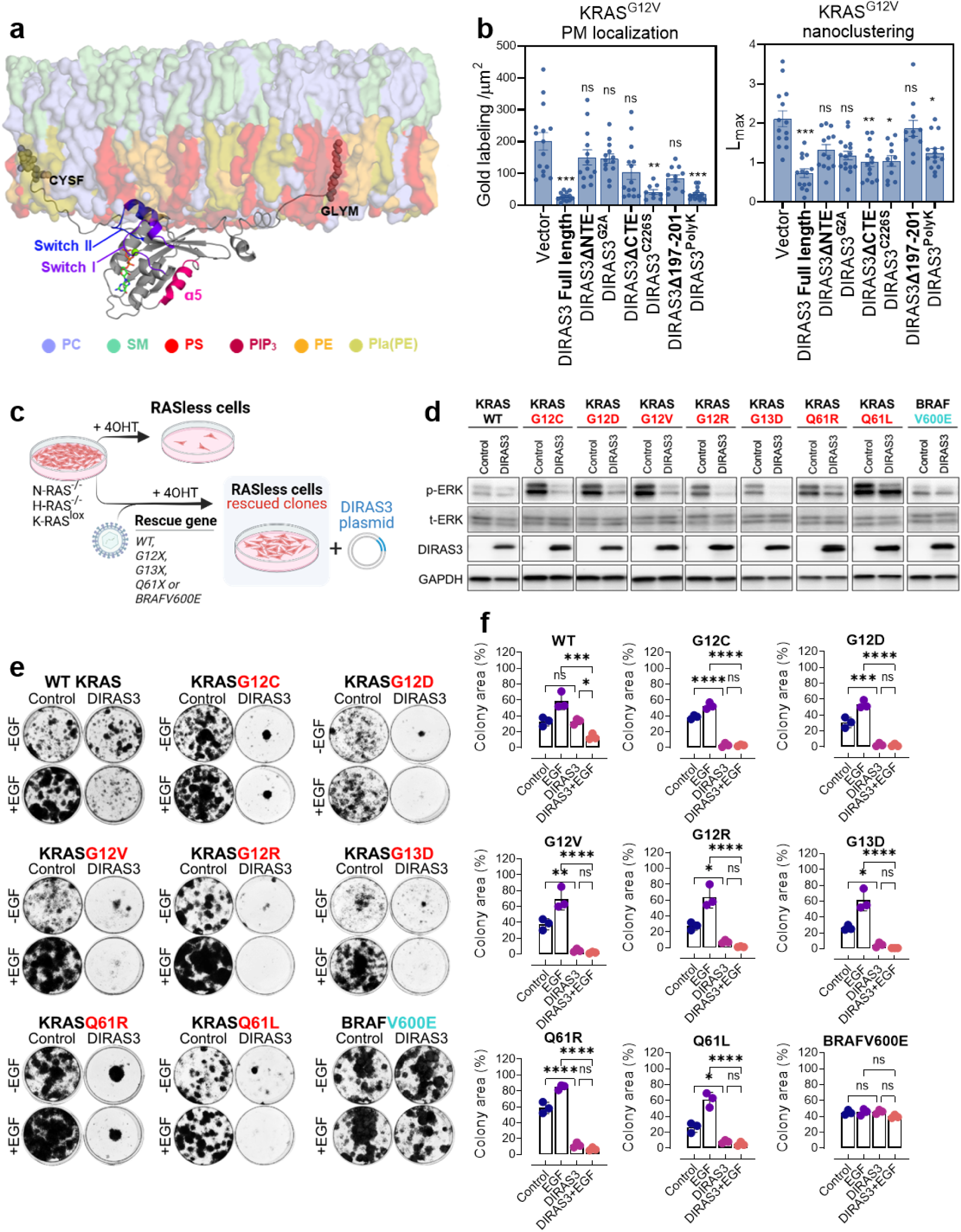
DIRAS3 α5 segment-dependent control of KRAS nanocluster organization and broad suppression of KRAS-dependent ERK activation and growth across oncogenic KRAS variants. **(a)** Representative snapshot from the molecular dynamics simulation showing the doubly anchored DIRAS3-plasma membrane complex. DIRAS3 is shown as a gray cartoon representation, with Switch I (purple), Switch II (blue), and the α5 helix (magenta) highlighted. **(b)** Electron microscopy analysis of KRAS nanoclustering following expression of WT or mutant DIRAS3. Basal PM sheets of cells expressing GFP-KRASG12V (GFP-KG12V) were transfected with empty vector, WT, or mutant DIRAS3 plasmids before being fixed and labeled with 4.5 nm gold anti-GFP before EM imaging. Univariate K-function (L(r)-r) analysis was performed to assess the spatial distribution of gold particles on the PM. Weighted mean univariate K-functions, the peak value of K-functions (Lmax), and mean gold labeling intensity for GFP-KG12V are shown. Values of L(r)-r above the 99% confidence interval (Cl) indicate significant clustering. Statistical significance of Lmax and gold labeling density was evaluated by bootstrap and two-tailed Student’s t-tests, respectively (mean ± SEM, n ≥ 12, ns p > 0.05; *p < 0.05; **p < 0.01; ***p < 0.001). **(c)** Schematic of the isogenic RASless MEF system reconstituted with WT KRAS, KRAS^G12X^, KRAS^G13X^, KRAS^Q61X^ alleles, or BRAF^V600E^. [Created in BioRender, https://BioRender.com/ol82yub]. **(d)** Representative immunoblot analysis of phospho-ERK1/2 and total ERK1/2 following transient expression of full-length DIRAS3 in RASless MEFs rescued with the indicated KRAS oncogenic alleles. Representative immunoblots from three independent biological experiments (n = 3) are shown. **(e)** Representative images of colony formation assays demonstrating reduced colony formation following transient expression of full-length DIRAS3 in KRAS-mutant RASless MEFs cultured in the presence or absence of EGF (20 ng/ml). Colonies were stained with Coomassie Brilliant Blue 10–14 days post-seeding. **(f)** Quantification of colony area relative to empty vector in KRAS-mutant RASless MEFs following transient expression of full-length DIRAS3 in the presence or absence of EGF (20 ng/mL). Representative triplicate images are shown in Supplementary Fig. 4A. Data are presented as mean ± SD from three independent biological experiments (*n* = 3). Statistical significance was determined by two-way ANOVA. ns, p > 0.05; ***p < 0.001; ****p < 0.0001.

Because KRAS nanoclustering at the plasma membrane is required for efficient effector recruitment and signal propagation^8, 15^, and MAPK output was tightly correlated with clonogenic growth across the DIRAS3 mutant panel, we determined whether differential suppression of ERK signaling reflected altered regulation of KRAS nanocluster organization. Plasma membrane sheets from cells expressing GFP-KRASG12V and doxycycline-inducible DIRAS3 were fixed and labeled with 4.5 nm gold-conjugated anti-GFP antibodies. Samples were imaged by electron microscopy and analyzed using linear K-functions, expressed as *L(r) - r*. Membrane localization of DIRAS3 was quantified by the gold labeling density (mm^2^), and the extent of nanoclustering was quantified by the maximum value (*L_max_*) of *L(r) – r*. The EM analysis confirmed robust plasma membrane localization of wild-type DIRAS3 upon dox treatment **(Supplementary Figure 3C-D)**. DIRAS3 itself formed nanoscale clusters at the plasma membrane **(Supplementary Figure 3E)**. The formation of DIRAS3 nanoclusters is consistent with that observed for several small GTPases within the RAS superfamily, including RAC1, RAP1, and KRAS^22, 23^. Wild-type DIRAS3 markedly disrupted the plasma membrane organization of KRAS, reducing both its membrane localization and nanoclustering. **(Figure 2B)**. In contrast, DIRAS3 mutants lacking the N-terminal extension or harboring the G2A mutation failed to disrupt KRAS nanoclusters, consistent with their complete loss of signaling and clonogenic growth inhibition. Mutants affecting the C-terminal CAAX motif or C226S prenylation exhibited partial impairment of nanocluster disruption **(Figure 2B)**, paralleling their intermediate effects on ERK phosphorylation and clonogenic growth **(Figure 1D-F)**.

Importantly, deletion of the α5-associated segment encompassing residues 197-201 abolished DIRAS3-mediated disruption of KRAS nanoclusters despite preserved membrane localization **(Figure 2B)**. This directly links the structural integrity of the DIRAS3 α5 helix, which is solvent-exposed and positioned for intermolecular interactions in the MD model **(Figure 2A)**, to the regulation of KRAS spatial organization. These findings demonstrate that membrane residence alone is insufficient for KRAS inhibition. Mutants retaining partial suppression of KRAS signaling and cell growth, including N197V and the poly-lysine cluster mutant, exhibited intermediate effects on nanocluster disruption, further reinforcing the relationship between spatial organization and signaling output.

Together, these data support a model in which DIRAS3 suppresses oncogenic KRAS signaling through topology-dependent membrane engagement coupled to α5-mediated interference with KRAS nanocluster formation or integrity. Dual lipid modification constrains DIRAS3 in a membrane-bound orientation that masks the canonical effector-binding interface while projecting the α5 helix towards the cytosol. This configuration enables DIRAS3 to perturb the KRAS plasma membrane interaction, reducing both membrane localization and nanoclustering, without directly engaging downstream effectors. Disassembly of KRAS nanoclusters attenuates MAPK signaling and constrains proliferative output, establishing a spatial mechanism for DIRAS3-mediated KRAS inhibition.

### DIRAS3 selectively targets active KRAS and functions as an endogenous RAS-ON pan-KRAS inhibitor

Having established the structural basis by which DIRAS3 engages KRAS and disrupts nanocluster-dependent signaling, we next investigated whether this inhibitory mechanism extends across oncogenic KRAS variants, including therapeutically challenging Q61 mutants. To address this question in a controlled genetic context, we employed RASless mouse embryonic fibroblasts (MEFs), which lack endogenous KRAS, HRAS, and NRAS and can be selectively rescued with individual oncogenic drivers. This system enables direct interrogation of KRAS-dependent signaling in an otherwise uniform cellular background^24^. RASless MEFs were rescued with wild-type KRAS, oncogenic KRAS alleles spanning the major hotspot codons (KRAS G12X, KRAS G13X, and KRAS Q61X), or the downstream effector mutation BRAF V600E **(Figure 2C)**.

Across all KRAS-mutant MEFs, expression of DIRAS3 produced a consistent inhibitory signature characterized by marked suppression of MAPK signaling, as indicated by reduced ERK1/2 phosphorylation, and significant impairment of clonogenic growth **(Figure 2D-F, Supplementary Figure 4A)**. These effects were observed across all tested oncogenic hotspot substitutions, demonstrating mutation-agnostic inhibition across the principal KRAS driver alleles examined. In contrast, DIRAS3 expression had minimal impact on MAPK signaling or clonogenic growth in MEFs expressing wild-type KRAS or BRAF-V600E under basal conditions, indicating that DIRAS3 impairs the oncogenic signaling of activated KRAS mutants.

To assess directly whether DIRAS3 activity correlates with KRAS activation, we stimulated wild-type KRAS MEFs with epidermal growth factor (EGF) to induce transient KRAS activation. Under these conditions, DIRAS3 exhibited robust inhibitory activity, significantly reducing clonogenic growth in EGF-stimulated wild-type KRAS cells **(Figure 2D-F, Supplementary Figure 4A)**. These findings indicate that DIRAS3 preferentially suppresses signaling in contexts where KRAS is activated and engaged in effector recruitment, consistent with its inhibition of oncogenic KRAS mutants.

Selectivity for activated KRAS was further supported by examining MEFs rescued with BRAF V600E, which signals downstream of KRAS independent of RAS activation. In this KRAS-independent context, DIRAS3 expression failed to suppress ERK1/2 phosphorylation or clonogenic growth, despite robust inhibition in KRAS-mutant cells **(Figure 2D-F, Supplementary Figure 4A)**. This selectivity demonstrates that DIRAS3 does not function as a general MAPK pathway inhibitor but instead acts through direct modulation of KRAS-dependent signaling assemblies. In addition to suppressing MAPK signaling and proliferation, DIRAS3 expression increased apoptotic cell death selectively in KRAS-dependent contexts (i.e., expressing oncogenic KRAS mutants or stimulated by EGF) **(Supplementary Figure 4B)**, further supporting its cytotoxic effects in cells reliant on oncogenic KRAS signaling.

Collectively, these data establish DIRAS3 as an endogenous antagonist of oncogenic KRAS that exhibits activation-biased, mutation-agnostic suppression across major hotspot alleles. Rather than targeting nucleotide cycling per se, DIRAS3 preferentially inhibits signaling-competent KRAS assemblies at the plasma membrane while sparing inactive or KRAS-independent signaling contexts. This spatially constrained mode of inhibition distinguishes DIRAS3 from nucleotide-state-targeting RAS-ON inhibitors and reveals a previously unrecognized endogenous mechanism for broad suppression of oncogenic KRAS signaling.

### A DIRAS3 α5-derived mimetic is sufficient to recapitulate activation-biased pan-KRAS inhibition

Our structural and functional analyses identified the DIRAS3 α5 helix as a critical determinant of KRAS engagement and nanocluster disruption, while the N- and C-terminal lipid modification establishes the membrane topology required for its proper spatial presentation. We therefore asked whether the α5 interface itself is sufficient to reproduce key aspects of KRAS inhibition, independent of the full-length DIRAS3 membrane scaffold. Demonstrating such sufficiency would indicate that the inhibitory activity of DIRAS3 is encoded within a discrete structural element, even though its physiological function is constrained by membrane-dependent orientation.

To test this, we evaluated a conformationally constrained cyclic DIRAS3-derived α5 mimetic peptide designed to recapitulate key structural determinants within the DIRAS3 α5 interface implicated in KRAS engagement **(Figure 3A and Supplementary Figure 5A-D)**^19^. The cyclic peptide spans the junction between the DIRAS3 α5 helix and hypervariable region (residues 194-203). As described previously, residues not predicted to contribute directly to KRAS binding were substituted with cysteines to enable thiol-based cyclization, and the N-terminal methionine was replaced with norleucine to improve oxidative stability^19^. The peptide incorporates critical α5 residues required for full-length DIRAS3-mediated KRAS inhibition and is cyclized to stabilize its bioactive helical conformation **(Figure 3A)**. An all-D-isomer version of the cyclic DIRAS3 peptide was synthesized as a control peptide for cell-based studies. As previously reported, the α5 peptide binds KRAS with mid-micromolar affinity, disrupts KRAS-RAF1 interactions, and exhibits efficient cytosolic access, enabling modulation of KRAS membrane organization^25, 26^. Structural modeling further supports a mechanism involving steric interference with the KRAS switch I interface required for RAF1 effector engagement **(Figure 3B)**.

**Figure 3.**
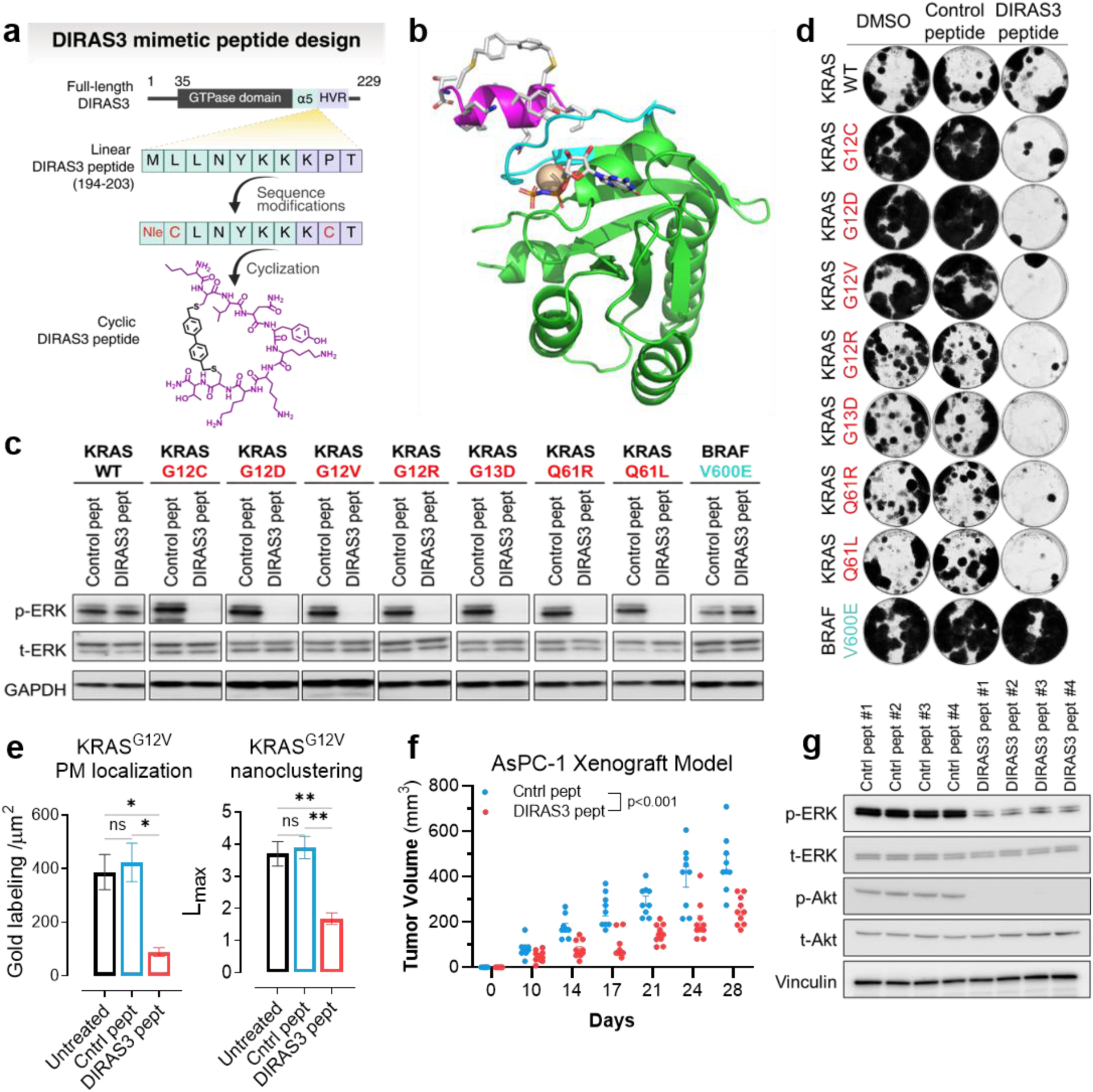
A DIRAS3-derived mimetic peptide recapitulates endogenous pan-KRAS inhibition. **(a)** Structure-guided design of a cyclic DIRAS3-derived mimetic peptide incorporating residues from the α5 interface and hypervariable region (HVR), including membrane-interacting motifs. The cyclization strategy is illustrated to enhance conformational stability. [Created in BioRender, https://BioRender.com/dy5l1tk]. **(b)** Structural model of the cyclic DIRAS3 mimetic peptide bound to KRAS. The peptide ribbon is magenta, with side chains colored by atom and N- and C-termini indicated. KRAS G12V (PDB 6GOE) is green, with switch I in cyan and GMPPNP and Mg^2+^ colored by atom. **(c)** Representative immunoblot analysis of phospho-ERK1/2 and total ERK1/2 in RASless MEFs rescued with the indicated KRAS^G12X^, KRAS^G13X^, and KRAS^Q61X^ alleles following treatment with DIRAS3 peptide or control peptide. Representative blots from three independent biological experiments (*n* = 3) are shown. **(d)** Representative colony formation assays showing reduced clonogenic growth in KRAS-mutant RASless MEFs treated with DIRAS3 peptide compared with control peptide. Colonies were stained with Coomassie Brilliant Blue 10– 14 d after seeding. Representative triplicate images are shown in Supplementary Fig. 6A, and quantification is shown in Supplementary Fig. 6B. **(e)** Quantitative electron microscopy image analysis of basal plasma membrane sheets from cells expressing GFP-KRAS^G12V^ following treatment with DIRAS3 peptide, control peptide, or untreated conditions. Gold labeling (4.5 nm anti-GFP) was used to visualize KRAS distribution. Univariate K-function (L(r)–r) analysis was used to assess the spatial clustering of gold particles. Weighted mean K-functions, peak clustering values (Lmax), and gold labeling density are shown. Values exceeding the 99% confidence interval indicate significant clustering. Statistical significance was determined by bootstrap analysis for Lmax and a two-tailed Student’s t-test for labeling density (mean ± SEM, n ≥ 12 membrane sheets; *p < 0.05; **p < 0.01). Representative K-function plots are shown in Supplementary Fig. 7B. **(f)** Evaluation of DIRAS3 peptide in an AsPC-1 pancreatic ductal adenocarcinoma xenograft model. Mice bearing subcutaneous AsPC-1 tumors were treated daily with DIRAS3 peptide (10 mg/kg) for 4 wk. Tumor growth was analyzed by two-way ANOVA. Each dot represents an individual tumor; error bars indicate SEM. **(g)** Representative immunoblot analysis of AsPC-1 xenograft tumors harvested at study endpoint (*n* = 4). Membranes were probed with the indicated antibodies.

We first assessed whether the DIRAS3 α5 mimetic peptide reproduces the mutation-agnostic inhibitory profile observed with full-length DIRAS3. Across RASless MEFs rescued with oncogenic KRAS alleles representing the major hotspot mutations (G12X, G13X, Q61X), treatment with the DIRAS3 peptide consistently suppressed MAPK signaling, as evidenced by reduced ERK1/2 phosphorylation, and significantly impaired clonogenic growth **(Figure 3C-D, Supplementary Figure 6A-B)**. This inhibitory pattern closely mirrored that observed following expression of full-length DIRAS3.

Consistent with the activation-state bias observed for full-length DIRAS3, the α5-derived peptide had minimal effects on MAPK signaling or growth in wild-type KRAS MEFs under basal conditions. However, upon epidermal growth factor stimulation, which transiently increases KRAS activation and membrane recruitment of RAF1, the peptide exhibited inhibitory activity in wild-type KRAS cells, reducing ERK phosphorylation and clonogenic growth **(Supplementary Figure 6C)**. These findings indicate that the isolated α5 region of DIRAS3 is sufficient to confer preferential inhibition of signaling-competent KRAS.

The specificity of the α5 mimetic for KRAS-dependent signaling was further confirmed in MEFs rescued with BRAF V600E. In this KRAS-independent context, the DIRAS3 peptide failed to suppress MAPK signaling or proliferation, paralleling the selectivity observed with full-length DIRAS3 **(Figure 3C-D, Supplementary Figure 6A-B)**. Thus, the α5-derived peptide recapitulates not only the inhibitory potency but also the pathway specificity of endogenous DIRAS3. Beyond MAPK suppression, treatment with the DIRAS3 α5 mimetic peptide increased apoptotic cell death in KRAS-mutant MEFs **(Supplementary Figure 7A)**, phenocopying the cytotoxic effects observed with full-length DIRAS3.

Mechanistically, the DIRAS3 α5 mimetic peptide perturbed signaling-competent KRAS membrane assemblies. Scanning electron microscopy analysis revealed reduced plasma membrane localization of KRAS G12V and marked disruption of KRAS nanocluster organization in peptide-treated cells compared with untreated or control peptide-treated cells **(Figure 3E, Supplementary Figure 7B)**. Confocal imaging of a fluorophore-conjugated peptide demonstrated detectable, but not predominant, plasma membrane localization **(Supplementary Figure 7C)**, with the majority of signal appearing as punctate intracellular structures. Thus, only a fraction of the peptide accesses the plasma membrane compartment. Given that RAF1 recruitment occurs selectively to GTP-bound KRAS within membrane nanoclusters, even limited peptide engagement at this interface may be sufficient to disrupt signaling-competent KRAS assemblies. Collectively, these findings suggest that the DIRAS3 α5 mimetic peptide suppresses MAPK signaling by perturbing KRAS nanocluster integrity, with disruption of effector engagement likely contributing to the destabilization of membrane-associated signaling assemblies, thereby recapitulating key features of the DIRAS3 inhibitory interface without requiring its full membrane-anchored topology.

To determine whether the KRAS-selective activity of the DIRAS3 peptide is maintained in broad genetically complex human cancers, we extend our analysis beyond the engineered RASless MEF system to a panel of pancreatic and ovarian cancer cell lines harboring either wild-type KRAS or distinct oncogenic KRAS hotspot mutations **(Supplementary Figure 8A-C)**. Despite the diverse mutation landscapes of these human cancer models, the DIRAS3 peptide consistently suppressed cell proliferation and induced apoptotic cell death across KRAS-mutant cancer cell lines, while exerting substantially weaker effects in KRAS wild-type cells **(Supplementary Figure 8A-B)**. These findings demonstrate that the anti-tumor activity of the DIRAS3 peptide is not restricted to a simplified genetic model but is preserved across genetically heterogeneous human cancers, underscoring its selective vulnerability toward oncogenic KRAS signaling and highlighting its therapeutic potential for KRAS-driven malignancies.

To determine whether the KRAS-inhibitory activity of the DIRAS3 peptide translated into therapeutic benefit in vivo, we evaluated its efficacy in an AsPC-1 PDAC xenograft model harboring oncogenic KRAS G12D. DIRAS3 peptide treatment significantly inhibited tumor growth relative to the control peptide group **(Figure 3F)**. Consistent with its mechanism of action in vitro, tumors from DIRAS3 peptide-treated mice exhibited reduced phosphorylation of ERK and AKT **(Figure 3G)**, supporting inhibition of KRAS-dependent MAPK and PI3K signaling in vivo. These findings demonstrate that pharmacologic mimicry of the DIRAS3 α5 domain effectively suppresses oncogenic KRAS signaling and tumor growth in vivo.

### DIRAS3 preserves dual MAPK and PI3K suppression in Q61-mutant contexts

Recently developed RAS-ON inhibitors have demonstrated substantial activity against multiple KRAS-driven tumors^27^. However, the efficacy of these agents can vary across KRAS alleles. We therefore investigated whether DIRAS3-mediated inhibition is maintained in Q61-mutant contexts, a clinically relevant subset that remains challenging for active-state-directed therapeutic strategies.

Q61-mutant KRAS proteins are characterized by impaired intrinsic and GAP-mediated GTP hydrolysis, resulting in sustained KRAS activation that is less dependent on upstream receptor tyrosine kinase input^28^. As a consequence, while RAS-ON inhibitors such as RMC-7977 effectively suppress MAPK signaling in G12-mutant contexts, Q61 mutants can maintain downstream signaling, particularly PI3K/AKT output, even when MAPK pathway activity is partially inhibited^28–30^. This mutation-specific signaling persistence presents a challenge for nucleotide-state-directed therapeutic strategies.

Given that DIRAS3 disrupts KRAS nanoclustering and effector engagement rather than targeting the nucleotide-binding pocket, we hypothesized that DIRAS3-mediated inhibition might retain pathway control in cells driven by Q61-mutant RAS. To test this directly, we compared the signaling and functional consequences of treatment with RMC-7977 and the DIRAS3 α5-derived mimetic peptide in RASless MEFs rescued with either KRAS G12D or KRAS Q61L.

In KRAS G12D and G12V MEFs, both RMC-7977 and the DIRAS3 peptide effectively suppressed MAPK signaling, as indicated by reduced ERK1/2 phosphorylation, and attenuated PI3K/AKT pathway activity **(Figure 4A)**. In contrast, in KRAS Q61L MEFs, RMC-7977 retained partial inhibition of MAPK signaling but failed to suppress PI3K/AKT activation, consistent with prior reports of differential pathway sensitivity in cells with RAS Q61-mutations. Under the same conditions, the DIRAS3 α5 peptide suppressed both MAPK and PI3K/AKT signaling outputs in KRAS Q61L cells, revealing a qualitative difference in pathway modulation between the two inhibitors.

**Figure 4.**
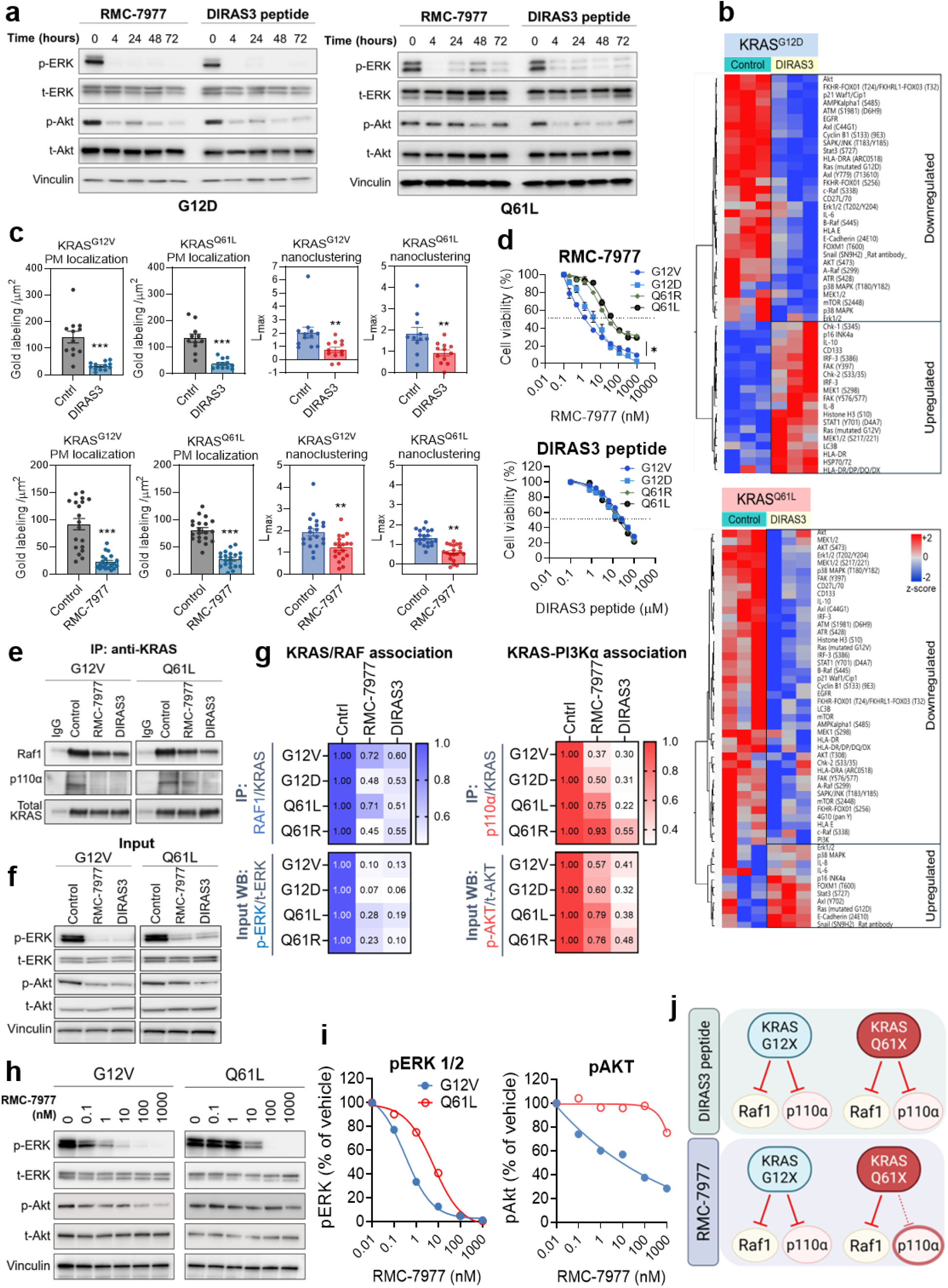
DIRAS3 mimetic peptide suppresses both MAPK and PI3K signaling across KRAS G12X and Q61X mutants. **(a)** Representative immunoblot analysis of MAPK (phospho-ERK1/2) and PI3K/AKT (phospho-AKT) signaling in RASless MEFs rescued with either KRAS^G12D^ or KRAS^Q61L^ alleles following treatment with RMC-7977 or the DIRAS3 peptide. Representative blots from three independent biological experiments (n = 3) are shown. **(b)** Reverse-phase protein array (RPPA) profiling of KRAS^G12D^ and KRAS^Q61L^ RASless MEFs following 15 μM control or DIRAS3 peptide treatment for 72h. Two-way hierarchical clustering was performed using Ward linkage and column standardization. Heatmap colors display relative protein expression levels as Z-scores, with red indicating higher and blue indicating lower expression. Data shown are mean-centered, normalized protein expression values from three independent biological replicates. **(c)** Quantitative electron microscopy analysis of basal plasma membrane sheets from KRAS^G12V^ and KRAS^Q61L^ cells treated with DIRAS3 peptide or RMC-7977. Gold-labeled GFP-KRAS was analyzed using univariate K-function (L(r)–r) analysis to assess spatial clustering. Weighted mean K-functions, peak clustering values (L_max_), and gold labeling density are shown. Values exceeding the 99% confidence interval indicate significant clustering. Statistical significance of Lmax was determined by bootstrap analysis and labeling density by two-tailed Student’s t-test (mean ± SEM, n ≥ 12 membrane sheets; ns, p > 0.05; **p < 0.01; ***p < 0.001). **(d)** Cell viability dose-response curves of KRAS^G12X^ and KRAS^Q61X^ RASless MEFs treated with DIRAS3 peptide or RMC-7977 for 72 h. Viability was measured using CellTiter-Glo and normalized to vehicle-treated controls. Data represent mean ± SD from three independent biological experiments (n=3). IC₅₀ values were calculated using nonlinear regression. Statistical analysis was performed using two-way ANOVA (ns, p > 0.05; *p < 0.05). **(e)** KRAS-effector interactions in RASless MEFs expressing KRASG12V or KRASQ61L following treatment with vehicle, RMC-7977 (10 nM), or DIRAS3 peptide (25 μM) for 4 hours. KRAS was immunoprecipitated from 500 μg of whole-cell lysates, and associated p110α and RAF1 were detected by immunoblotting; immunoprecipitated KRAS served as the loading control. Blots are representative of 2 independent biological replicates. **(f)** Immunoblots of the corresponding input lysates from the experiments in (e) and Supplementary Figure 8E-F, probed for phospho-ERK1/2, total ERK1/2, phospho-AKT, total AKT, and Vinculin. Loss of RAS– RAF1 association in (e), together with reduced ERK phosphorylation here, confirms target engagement by RMC-7977 at the concentration used. Blots are representative of 2 independent biological replicates. **(g)** Quantification of the blots in (e) and (f). Band intensities were measured by densitometry in ImageJ. Immunoprecipitated RAF1 and p110α were normalized to total RAS recovered in the same immunoprecipitate; input phosphoproteins were normalized to their respective total protein. Values are expressed as fold change relative to vehicle-treated cells of the same genotype and displayed as a heatmap, with darker colors indicating greater retention of signal and lighter colors indicating loss relative to vehicle. Blots for G12D and Q61R are shown in SF-8E-F. **(h)** Dose-dependent effects of RMC-7977 on MAPK and PI3K-AKT signaling. RASless MEFs expressing KRASG12V or KRASQ61L were treated for 4 h with DMSO vehicle or RMC-7977 at 0.1, 1, 10, 100, or 1000 nM. Whole-cell lysates were immunoblotted for phospho-ERK1/2, total ERK1/2, phospho-AKT, total AKT, and vinculin. Blots are representative of 3 independent biological replicates. **(i)** Quantification of the blots in (h). Band intensities were measured by densitometry in ImageJ; phospho-signals were normalized to the corresponding total protein and expressed as a percentage of vehicle-treated cells of the same genotype. Vehicle is plotted at 0.01 nM to permit display on a logarithmic axis. At 100 nM RMC-7977, a concentration producing ≥95% suppression of ERK phosphorylation in both lines, p-AKT was 37 % of vehicle in KRASG12V cells and 97% in KRASQ61L cells. **(j)** Proposed model for the differential effects of RMC-7977 and DIRAS3 peptide in KRASG12X and KRASQ61X cells. RMC-7977 displaces RAF1 but not p110α from Q61-mutant KRAS, whereas the α5 peptide occludes both effector interfaces independently of nucleotide state. [Created in BioRender, https://BioRender.com/prxhci1].

To globally assess pathway regulation, we performed reverse-phase protein array (RPPA) profiling of KRAS G12D and KRAS Q61L MEFs treated with the DIRAS3 peptide (versus treated with control peptide). This analysis revealed concordant downregulation of several MAPK and PI3K pathway components across both mutant backgrounds upon DIRAS3 peptide treatment **(Figure 4B)**, reinforcing the mutation-spanning inhibitory profile of DIRAS3.

Mechanistically, both DIRAS3 and RMC-7977 disrupted KRAS nanoclustering across G12- and Q61-mutant backgrounds **(Figure 4C)**, as confirmed by scanning electron microscopy, which revealed comparable perturbation of KRAS membrane nanoclusters. Despite these similar effects on membrane organization, downstream signaling outputs diverged in KRAS Q61-mutant cells. Whereas both agents suppressed MAPK signaling, only DIRAS3 maintained coordinated suppression of both MAPK and PI3K signaling, consistent with the activity of the DIRAS3 α5 mimetic. Aligned with dual-pathway inhibition, KRAS Q61L MEFs remained sensitive to DIRAS3-mediated growth suppression, despite exhibiting reduced responsiveness to RMC-7977 **(Figure 4D)**. DIRAS3 induced comparable growth inhibition in Q61- and G12-mutant cells, indicating robust activity across KRAS alleles with differential sensitivity to RAS-ON inhibitors.

To determine whether these differences reflected distinct effects on KRAS-effector interactions, we performed KRAS co-immunoprecipitation in RASless MEFs expressing KRASG12V, KRASG12D, KRASQ61L, or KRASQ61R. In KRASG12X cells, both RMC-7977 and the DIRAS3 peptide reduced the association of KRAS with p110α and RAF1, accompanied by decreased AKT and ERK phosphorylation **(Figure 4E-G, Supplementary Figure 8E-F)**. In KRASQ61X cells, both agents disrupted KRAS-RAF1 association and reduced ERK phosphorylation. However, RMC-7977 had little effect on KRAS-p110α association or AKT phosphorylation, whereas the DIRAS3 peptide suppressed both **(Figure 4E-G, Supplementary Figure 8E-F)**. To test whether the retention of AKT signaling in Q61-mutant cells reflected incomplete target engagement, we treated KRASG12V and KRASQ61L RASless MEFs with RMC-7977 across a four-log range (0.1– 1000 nM) for 4 h **(Figure 4H-I)**. ERK phosphorylation was abolished in both lines at ≥100 nM, although Q61L cells were less sensitive at lower concentrations. In contrast, AKT phosphorylation showed a marked genotype-dependent difference: at 100 nM, a concentration producing ≥95% ERK suppression in both genotypes, AKT phosphorylation remained 37% of vehicle levels in G12V cells and 97% in Q61L cells **(Figure 4H-I)**. Preservation of AKT phosphorylation despite near-complete ERK suppression in the same Q61L lysates rules out insufficient drug exposure and supports differential suppression of the two signaling pathways. These findings indicate that although RMC-7977 and the DIRAS3 peptide similarly suppress RAF1-ERK signaling, the DIRAS3 peptide more broadly disrupts KRAS effector engagement, particularly PI3K-AKT signaling driven by KRASQ61L **(Figure 4J)**. Thus, although RMC-7977 and the DIRAS3 peptide similarly suppress RAF1-ERK signaling in Q61L cells, only the DIRAS3 peptide also disrupts KRAS-p110α association and PI3K-AKT signaling.

Together, these data reveal a fundamental mechanistic distinction between nucleotide-state targeting RAS-ON inhibitors and DIRAS3-mediated KRAS inhibition. Whereas RMC-7977 exhibits mutation-dependent differences in suppression of the PI3K-AKT pathway, DIRAS3 preserves coordinated inhibition of MAPK and PI3K signaling across multiple oncogenic KRAS mutations, including Q61 mutants. These findings support a model in which DIRAS3 combines disruption of nanocluster organization with direct interference with KRAS-effector engagement, providing a mechanistically distinct strategy that may overcome mutation-dependent limitations of nucleotide-state-directed inhibition.

## Discussion

In this study, we define the structural and spatial basis by which DIRAS3 suppresses oncogenic KRAS signaling and establish a mechanistically distinct paradigm for pan-mutant KRAS inhibition. We demonstrate that DIRAS3-mediated suppression depends on topology-dependent membrane engagement mediated by dual lipid anchoring to an α5-encoded interface that disrupts signaling-competent KRAS assemblies. Collectively, our findings support a model in which spatial perturbation of KRAS nanocluster organization, coupled with disruption of effector engagement, enables the mutation-spanning inhibitory activity of DIRAS3.

We first delineated the structural determinants required for KRAS engagement and inhibition. Dual lipid modification, N-terminal myristoylation and C-terminal prenylation, was essential for maximal activity, with N-terminal anchoring contributing disproportionately to effective KRAS suppression. Unlike canonical RAS proteins, which rely primarily on C-terminal CAAX-mediated prenylation for membrane association, DIRAS3 requires coordinated lipid N-terminal and C-terminal anchoring to establish the membrane topology necessary for function. Molecular dynamics-based membrane topology analysis further supported a membrane orientation in which the α5 region remains solvent-exposed while the canonical switch-region effector interface is positioned toward the membrane. Importantly, deletion of the α5-associated segment (residues 197-201) abolished KRAS inhibition despite preserved membrane localization, directly linking this structural element to disruption of KRAS spatial organization. These findings indicate that membrane residence of DIRAS3 alone is insufficient; rather, proper membrane orientation imposed by dual lipid anchoring enables α5-mediated engagement required to interfere with KRAS nanocluster integrity.

A central mechanistic insight from this work is that KRAS nanocluster organization and effector engagement are closely linked processes that together sustain signaling output. In the context of full-length DIRAS3, dual lipid anchoring constrains the protein within the plasma membrane, positioning the α5 interface to engage KRAS within signaling-competent assemblies. In contrast, the DIRAS3 α5 mimetic peptide lacks membrane-anchoring elements and therefore does not recapitulate this topology. Instead, it functions through a distinct, topology-independent mechanism, directly interfering with effector engagement.

Recruitment of RAF1 to the plasma membrane occurs preferentially upon KRAS GTP loading, when active KRAS assembles into nanoclusters that function as signaling platforms. Effector engagement is not merely a consequence of clustering; rather, RAF1 binding stabilizes and reinforces signaling-competent KRAS assemblies. Recent studies further support the concept that KRAS-effector interactions exist within a dynamic competitive equilibrium in which stabilization of RAF1-containing assemblies can oppose inhibitory perturbations targeting active KRAS complexes ^4^. Within this framework, our findings suggest that full-length DIRAS3 perturbs the spatial organization of signaling-competent KRAS nanoclusters through topology-dependent membrane engagement imposed by dual lipid anchoring. Although we did not directly examine RAF1 displacement by full-length DIRAS3, disruption of nanocluster architecture would be expected to impair the stability of effector-associated signaling assemblies.

In contrast, the isolated DIRAS3 α5 mimetic peptide functions through a distinct topology-independent mechanism. Structural modeling predicts that the α5 mimetic occupies a surface positioned to interfere directly with KRAS-RAF1 engagement **(Figure 3B)**, consistent with prior NMR-based biophysical evidence demonstrating disruption of KRAS-RAF1 interactions^19^. Within this context, interference with effector engagement likely contributes to the coordinated loss of clustering and downstream MAPK and PI3K signaling observed in our system.

Consistent with this spatial mechanism, the sufficiency of the α5 helix to recapitulate key functional features of full-length DIRAS3 establishes that the inhibitory mechanism is structurally encoded within a discrete module. The conformationally constrained α5 mimetic peptide reproduced mutation-agnostic suppression across G12, G13, and Q61 hotspot alleles, retained activation bias, disrupted KRAS nanoclusters, blocked RAF1 engagement, and induced KRAS-dependent cytotoxicity. This activity is achieved in the absence of membrane-anchored topology, indicating that while full-length DIRAS3 relies on membrane-imposed topology to position the α5 interface, the isolated α5 segment can function independently by directly targeting effector interactions. This distinction highlights two mechanistically related but operationally distinct modes of KRAS inhibition. Together, these findings suggest that the α5 interface encodes a core inhibitory module that can be deployed through distinct mechanistic architectures. In the context of full-length DIRAS3, membrane topology imposed by dual lipid anchoring enables spatial perturbation of signaling-competent KRAS assemblies. In contrast, the isolated α5 mimetic peptide functions independently of membrane-constrained organization and directly interferes with KRAS-effector engagement Importantly, our findings also reveal important mechanistic distinctions between DIRAS3-mediated inhibition and recently developed active-state RAS inhibitors ^4, 27, 31^. Emerging evidence indicates that RAS-ON inhibitors function not solely through nucleotide-state recognition but also through the formation of inhibitor-induced complexes that sterically restrict effector access to GTP-bound KRAS and destabilize signaling-competent RAS assemblies. Recent studies further suggest that disruption of these higher-order KRAS-effector complexes contributes to their therapeutic activity and may influence resistance mechanisms in RAS-driven tumors^4^. In this setting, both DIRAS3-derived inhibition and RAS-ON inhibitors converge on suppression of effector-coupled KRAS signaling platforms, although the mechanisms by which these perturbations are achieved differ.

Whereas tri-complex RAS inhibitors such as daraxonrasib rely on the formation of drug-induced inhibitory interfaces that competitively restrict effector engagement with active KRAS^4^, full-length DIRAS3 appears to function primarily through topology-dependent perturbation of KRAS spatial organization imposed by dual lipid anchoring. The isolated α5 mimetic peptide, by contrast, directly interferes with KRAS-effector engagement independently of membrane-constrained topology. These mechanistic distinctions may be particularly relevant to Q61-mutant KRAS proteins, which exhibit altered switch-II dynamics and differential pathway dependencies. In our system, both RMC-7977 and the DIRAS3 α5 mimetic disrupted KRAS nanocluster organization in G12- and Q61-mutant contexts, indicating that nanocluster perturbation alone is insufficient to explain their differential signaling effects. Although RMC-7977 retained suppression of RAF1-ERK signaling in Q61-mutant cells, it showed limited inhibition of KRAS-p110α association and PI3K-AKT signaling. In contrast, the DIRAS3 α5 mimetic disrupted association of KRAS with both RAF1 and p110α and preserved coordinated suppression of MAPK and PI3K signaling across G12- and Q61-mutant backgrounds. These findings suggest that DIRAS3 and RMC-7977 can similarly perturb KRAS membrane organization while differing in their ability to restrict specific interactions. Prior NMR analysis of the DIRAS3 α5 interface supports direct interaction with KRAS regions involved in effector engagement, including the switch-I region associated with RAF1 binding, with weaker effects on adjacent interfaces, including switch-II^19^. The ability of the DIRAS3 peptide to disrupt p110α association with Q61L KRAS raises the possibility that its direct engagement of KRAS can interfere with PI3K-effector interactions that are relatively less sensitive to RMC-7977 treatment. Thus, direct engagement of KRAS by the DIRAS3 peptide may provide a mechanistic route to broader effector inhibition that is less sensitive to mutation-dependent differences in the response to nucleotide-state-directed inhibitors. Structural studies will be required to define the precise basis of this differential effector inhibition. More broadly, these findings suggest that targeting both the spatial organization and effector accessibility of KRAS signaling assemblies may provide a mechanistically distinct approach to suppressing oncogenic KRAS across mutation classes.

Several limitations of this study should be considered. First, much of our mechanistic analysis was conducted in RASless MEFs reconstituted with defined oncogenic drivers, which enables precise interrogation of KRAS-dependent signaling, but does not fully reflect the complexity of endogenous tumor signaling involving multiple RAS isoforms. Although we demonstrated anti-tumor activity of the DIRAS3 peptide in an AsPC-1 PDAC xenograft model and previously reported that full-length DIRAS3 suppresses tumor growth in pancreatic cancer models harboring G12D or G12V KRAS mutations ^10, 19, 32^, broader evaluation across genetically diverse tumor models and additional in vivo systems will be important to define the generalizability and physiological relevance of this mechanism. Second, although the α5-derived mimetic reproduces key functional features of full-length DIRAS3, peptide delivery and intracellular distribution remain suboptimal. Further optimization of stability, membrane targeting, and pharmacologic properties will be necessary to enhance translational potential. Finally, the long-term adaptive consequences of spatial KRAS inhibition remain unclear, including the potential for compensatory rewiring of parallel signaling pathways, which can confer resistance to other modes of KRAS inhibition. Addressing these issues will help clarify the therapeutic feasibility of effector-coupled spatial interference as a strategy for RAS-driven cancers.

Together, our study identifies DIRAS3 as an endogenous, activation-biased pan-KRAS inhibitor and establishes perturbation of effector-coupled KRAS signaling assemblies as a distinct mechanism for broadly suppressing oncogenic KRAS signaling. We further demonstrate that the inhibitory activity encoded within the DIRAS3 α5 interface can be deployed through mechanistically distinct architectures, including topology-dependent spatial regulation in the context of full-length DIRAS3 and direct effector interference by an isolated α5-derived mimetic peptide. The preferential activity of DIRAS3 toward oncogenic versus wild-type KRAS suggests that endogenous regulation of KRAS signaling assemblies may enable selective suppression of mutant KRAS signaling. Consistent with this model, the α5 mimetic suppresses signaling across multiple KRAS variants, including the Q61 mutant that remains challenging for current RAS-ON inhibitor strategies, supporting the concept that higher-order KRAS signaling organization can be therapeutically targeted independently of nucleotide-state selectivity. More broadly, these findings identify naturally evolved inhibitory interfaces as a tractable source of therapeutic strategies and reveal the spatial organization of oncogenic KRAS assemblies as an underappreciated vulnerability in KRAS-driven cancers.

## Materials and Methods

### RASless MEFs

RASless MEFs were generated as previously described by Barbacid and colleagues^1,2^ and provided by the Frederick National Laboratory of the National Cancer Institute Ras Initiative/Leidos Biomedical Research. Briefly, MEFs were isolated from embryos derived from mice lacking Hras and Nras alleles and harboring a conditional Kras allele (Kras^lox/lox^) together with a tamoxifen-inducible Cre recombinase (Cre-ERT2). Following establishment of primary MEF cultures, cells were treated with 4-hydroxytamoxifen (4-OHT) to induce Cre-mediated excision of the floxed Kras allele, thereby eliminating expression of all three endogenous RAS isoforms. Complete ablation of RAS results in proliferative arrest, which provides a genetically defined system to investigate the functions of exogenously introduced KRAS variants (WT, G12C, G12V, G12D, G13D, G12R, Q61R, Q61L) or downstream signaling components (BRAFV600E) in the absence of confounding endogenous RAS activity.

### Cell culture and maintenance

RASless MEF clones were maintained at 37°C in a humidified atmosphere with 5% CO₂. Cells were cultured in Dulbecco’s modified Eagle’s medium (DMEM) supplemented with 10% fetal bovine serum (FBS) and 1% penicillin-streptomycin. Cells were routinely passaged at ∼70–80% confluence using 0.05% trypsin-EDTA. The experiments were conducted in early passages, typically between passages 2 and 8, and never beyond passage 9 to ensure optimal viability and reproducibility. Pancreatic ductal adenocarcinoma cell lines were obtained from ATCC including AsPC-1 (KRASG12D; CRL-1682), BxPC-3 (KRAS WT, BRAFV600E; CRL-1687), Capan-2 (KRASG12V; HTB-80), and MIA PaCa-2 (KRASG12C; CRL-1420). The low-grade serous ovarian cancer cell line HCC5075 (KRASG12V) was obtained from Dr. Adi F. Gazdar, UT Southwestern Medical Center (Dallas, TX). VOA-3448 (KRAS WT) cells were obtained from Dr. Mark S. Carey, University of British Columbia (Vancouver, Canada). PM-LGSOC-01 (KRASG12V) cells were provided by Dr. Olivier De Wever, Ghent University (Ghent, Belgium). The OVCAR-8 (KRASP121H) and Hey (KRASG12D) ovarian cancer cells were provided by Dr. Gordon Mills, UT M.D. Anderson Cancer Center (Houston, TX). AsPC-1, Hey, BxPC-3, MIA PaCa-2, OVCAR-8, and HCC5075 cells were routinely grown in RPMI1640 (Corning, 15–040-CV) supplemented with 2 mM L-glutamine, and 10% fetal bovine serum (FBS), 100 U of penicillin and 100 μg of streptomycin. Capan-2 cells were grown in McCoy’s (Corning, 10–050-CV) supplemented with 10% FBS, 100 U of penicillin, and 100 μg of streptomycin. PM-LGSOC-01 cells were grown in EMEM (Corning, 15–010-CV) supplemented with 10% FBS, 100 U of penicillin, and 100 μg of streptomycin. VOA-3448 cells were grown in 1:1 mixture of M199 and MCDB media supplemented with 10% FBS, 100 U of penicillin, and 100 μg of streptomycin. All cells were maintained in a humidified atmosphere of 5% CO2 and 95% room air.

### Transfections

Individual RASless MEF clones were maintained at 37 °C in Dulbecco’s modified Eagle’s medium supplemented with 10% fetal bovine serum. Cells were seeded 24 h prior to transfection to achieve 60–70% confluency at the time of transfection. For transient expression, cells were transfected with 1 µg of pcDNA3.1(+) plasmids encoding wild-type or mutant DIRAS3 constructs using FuGENE HD (Promega) at a 3:1 reagent-to-DNA ratio, according to the manufacturer’s instructions in antibiotic-free medium. Transfection efficiency and expression of DIRAS3 constructs were verified 24 h post-transfection by immunoblot analysis of whole-cell lysates using anti-DIRAS3 antibody. Robust overexpression relative to empty vector-transfected controls was consistently observed across experiments. Cells were incubated for 24-72 h post-transfection at 37°C, depending on the downstream assay (e.g., immunoblotting, colony formation, or signaling analyses), and subsequently harvested for analysis.

### Colony formation assay

Cells were trypsinized, counted, and seeded at a density of 500 cells per well in 6-well or 12-well plates in triplicate. Cells were cultured for 10-14 days to allow colony formation, with medium replaced every 3-4 days. At the endpoint, colonies were gently washed with PBS and stained with 0.1% Coomassie Brilliant Blue solution (0.1% Coomassie Brilliant Blue, 30% methanol, 10% acetic acid). Plates were rinsed thoroughly with distilled water and air-dried. Colony growth was quantified by measuring total colony area per well using ImageJ. Images were captured under identical settings, converted to 8-bit, background-subtracted, and thresholded uniformly across all samples. The cumulative colony area per well was calculated and used for statistical analysis. Data represent the mean ± SEM from three independent experiments.

### Immunoblotting

To extract protein from cells, the wells were washed with cold PBS and treated with SDS lysis buffer for 1 min at room temperature. The lysis buffer contained Tris-HCl (pH 6.8), SDS, glycerol, β-mercaptoethanol, and bromophenol blue. The lysates were heated at 99°C for 5 min and then separated using SDS-PAGE with a 5-12% polyacrylamide gel. The separated proteins were transferred to PVDF membranes and subjected to immunoblot analysis with the specified antibodies. The protein bands were visualized using an ECL-enhanced chemiluminescence detection kit (SuperSignal™ West Pico PLUS, Thermo Scientific). The intensity of the bands in the Western blots was quantified using the ImageJ program.

### Antibodies

Antibodies against phospho-ERK^Thr202/Tyr204^ (Cat# 4370), ERK (Cat# 9102), Phospho-Akt^Ser473^ (Cat# 4060), Akt (Cat# 9272), GAPDH (Cat# 2118), RAF1 (#53745), and p110α (#4249) were purchased from Cell Signaling Technology. Anti-Vinculin (V9264) was purchased from Sigma-Aldrich. The anti-DIRAS3 antibody was generated and characterized by our group.

### Generation of human PDAC xenografts in mice

For the DIRAS3-inducible mouse study evaluating early signaling changes, DIRAS3-inducible or parental PDAC AsPC-1 cells (1.5 × 10^6^) were injected subcutaneously into eight-week-old male athymic nu/nu mice (Envigo Biosciences). Each group contained five mice. To assess early signaling changes in vivo, doxycycline (DOX) administration was initiated one week after tumor cell injection and continued for 8 days. Mice received drinking water containing 2 mg/mL DOX in 5% sucrose (DIRAS3+) or 5% sucrose alone (DIRAS3−). Water was replaced every two days throughout the study. Tumors were harvested at the end of the 8-day induction period for analysis of MAPK pathway activity, including phospho-ERK levels. For the AsPC-1 xenograft model used for peptide treatment studies, mice were injected subcutaneously in the right flank with AsPC-1 (1.5 × 10^6^) cells harboring a KRASG12D mutation. Once tumors became palpable, animals were randomized into two groups (n = 10 per group) and treated intraperitoneally with 10 mg/kg control or DIRAS3 peptide. Treatment continued daily for four weeks. Tumor volume was measured every three days using calipers. Statistical analyses were performed in GraphPad Prism using two-way ANOVA to assess the effects of peptide treatment on tumor growth. All animal experiments were conducted in accordance with institutional and national guidelines for the care and use of laboratory animals and were approved by the Institutional Animal Care and Use Committee (IACUC) of The University of Texas MD Anderson Cancer Center under protocol number 00001195-RN03.

### Mass spectrometry

Harvested cells were lysed by sonication in 2.5 volumes of lysis buffer containing 25 mM Tris (pH 7.5), 1 mM EDTA, 0.5% NP-40, 0.1% CHAPS, and 150 mM KCl. Lysates were cleared by centrifugation at 100,000 × g, and the supernatants were subjected to immunopurification using an in-house anti-DIRAS3 polyclonal antibody. Affinity-purified DIRAS3 proteins were digested on beads. Briefly, beads were washed twice with cold PBS, resuspended, and digested with 500 ng trypsin (T9600, GenDepot) at 37°C for 3 h. Digested peptides were enriched using in-house STAGE tips packed with 2 mg C18 beads^33^ (3 µm, Dr. Maisch GmbH, Germany) and dried under vacuum. Peptides were resuspended and analyzed using an EASY-nLC 1000 system (Thermo Scientific) coupled to an Orbitrap Fusion mass spectrometer (Thermo Scientific) equipped with an ESI source. Peptides were loaded onto an in-house ReproSil-Pur Basic C18 trap column (1.9 µm, Dr. Maisch GmbH, Germany; 2 cm length, 100 µm i.d.) and separated on a 5 cm analytical column (150 µm i.d.) using a 75 min gradient of 0–35% acetonitrile/0.1% formic acid at a flow rate of 800 nL/min. Data acquisition was performed using both data-dependent acquisition (DDA) mode for unbiased peptide identification and parallel reaction monitoring (PRM) mode for targeted detection of the myristoylated N-terminal peptide. For DDA analysis, precursor MS spectra were acquired over a mass range of 300–1400 m/z at a resolution of 120,000 (at 400 m/z) using the Orbitrap analyzer with an AGC target of 7.5 × 10^5^ and a maximum injection time of 50 ms. A top-speed method with a 2 s cycle time was used for precursor selection. Selected MS1 ions were isolated using the quadrupole with a 1.6 m/z isolation window and 15 s dynamic exclusion, fragmented by higher-energy collisional dissociation (HCD), and analyzed in the ion trap using rapid scan mode with an AGC target of 5 × 10^3^ and a maximum injection time of 35 ms. For PRM analysis of N-terminal glycine myristoylation, the targeted precursor ion was m/z 489.2914 (2+ charge state; myr-GNASFGSK). The precursor ion was monitored throughout the 75 min LC-MS run and isolated by quadrupole, followed by collision-induced dissociation (CID)-based MS/MS analysis. Acquired spectra were searched against the target-decoy Human RefSeq database (release 2020) using Proteome Discoverer version 2.1 (Thermo Fisher Scientific) with the Mascot search engine (version 2.4, Matrix Science). Dynamic modifications included N-terminal glycine myristoylation, protein N-terminal acetylation, and methionine oxidation. Precursor and fragment mass tolerances were set to 20 ppm and 0.5 Da, respectively, with up to two missed tryptic cleavages allowed. Peptide-spectrum matches were filtered to a 1% false discovery rate (FDR) using Percolator-based validation.

### Cell viability assay

Cells were seeded at a density of 5 × 10^3^ cells per well in black-walled, clear-bottom 96-well plates and allowed to attach overnight at 37°C in a humidified atmosphere containing 5% CO_2_. The following day, culture medium was replaced with fresh medium containing serial dilutions of the indicated compounds. Cells were incubated for 72 h under standard culture conditions. Cell viability was assessed using the CellTiter-Glo® 2.0 assay (Promega) according to the manufacturer’s protocol. Briefly, an equal volume of CellTiter-Glo reagent (100 µl) was added to each well. Plates were shaken for 2 min to induce cell lysis and incubated for 10 min at room temperature to stabilize the luminescent signal. Luminescence was measured using a BioTek microplate reader. Background luminescence from wells containing medium and reagent only was subtracted. Relative luminescence units (RLU) were normalized to vehicle-treated controls, and cell viability was expressed as a percentage of control. Experiments were performed in triplicate wells and repeated in at least three independent biological replicates.

### Caspase 3/7 Glo assay

Caspase-3/7 activity was measured using the Caspase-Glo® 3/7 Assay kit (Promega) following the manufacturer’s instructions. Briefly, RASless MEFs were seeded at 5 × 10³ cells per well in black-walled 96-well plates and allowed to attach overnight. Following the indicated treatments, an equal volume of Caspase-Glo reagent was added directly to each well. Plates were gently mixed and incubated for 30 min at room temperature in the dark. Luminescence was recorded using a BioTek microplate reader. Background values from wells containing medium and reagent only were subtracted. Caspase activity was expressed as fold change relative to vehicle-treated controls. Experiments were performed in triplicate wells and repeated in at least three independent biological replicates.

### Computational analysis of DIRAS3 membrane topology

The full-length, GTP-bound structure of DIRAS3 was modeled using AlphaFold3^34^, and the protein-membrane complex was constructed using CHARMM-GUI^35, 36^. The N-termini Gly2 was patched with a myristoyl chain, and a farnesyl isoprenoid lipid was added to the C-terminal Cys226. The protein was then attached to an asymmetric bilayer modeling the plasma membrane (PM) and made up of equal numbers of phosphatidylcholine (PC) and sphingomyelin (SM) in one leaflet and PC, phosphatidylserine (PS), phosphatidylinositol (3,4,5)-trisphosphate (PIP3), phosphatidylethanolamine (PE), and plasmalogen-PE (Pla(PE)) in the opposing leaflet where DIRAS3 is attached. The leaflet representing the inner monolayer of the PM contained ∼3% molar concentration of PIP3, and the remaining lipids, including PC, PS, PE, and Pla (PE) were distributed equally, following previous work on DIRAS3 N-terminal extension^37^. Simulations were conducted using the CHARMM36m force field and TIP3P waters containing 150 mM of Na+ and Cl-ions. All-atom molecular dynamics (MD) simulations were performed on the ANTON 3 for 20 µs after initial equilibration on local resources using GROMACS^38, 39^.

### Scanning Electron Microscopy Assay of KRAS Nanoclustering

DIRAS3 expression was achieved by transfecting cells as described above in the transfection methods. Both DIRAS3-negative and DIRAS3-positive cells were then infected with lentiviral GFP-KRAS^G12V^ or GFP-KRAS^Q61L^ 18 h prior to harvesting. EM grids and adherent cells were collected for scanning electron microscopy analysis of KRAS nanoclustering. Briefly, the basolateral membrane was exposed and fixed using 4% paraformaldehyde (PFA) and 0.1% glutaraldehyde (GA). For the univariate experiments, 4.5-nm-gold-anti-GFP antibodies were used to immunolabel GFP-RAS on the PM and embedded in uranyl acetate. Gold particle distribution was visualized by transmission EM (JEOL JEM-1400). ImageJ (RRID:SCR_003070) was then used to assign x and y coordinates of gold particles in a 1-µm^2^ area on an intact and featureless PM region. Ripley’s K-Function was used to calculate the gold particle distribution and extent of nanoclustering. (Equations A and B):

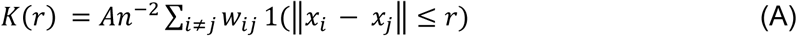

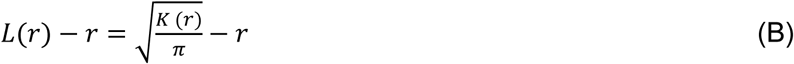

where *K*(*r*) = the univariate K-function for a pattern of *n* points in a selected PM area of A; r = length scale with the range of 1 < r < 240 nm evaluated at increments of 1 nm; ‖ · ‖ is Euclidean distance; 1(·) is the indicator function with a value of 1 if ‖*x_i_* − *x_j_*‖ ≤ *r*, and a value of 0 otherwise; and *w_ij_^-1^* is the fraction of the circumference of a circle with center *x_i_* and radius ‖*x_i_* − *x_j_*‖ contained within area *A*, rendering an unbiased edge correction for points at the edge of the study area. *K(r)* is transformed into *L(r) – r,* which is then normalized on the 99% confidence interval (99% C.I.) estimated via Monte Carlo simulations. Under the null hypothesis of complete spatial randomness, *L(r) – r =* 0 for all values of *r*. Values of *L(r) – r* exceeding the confidence interval indicate significant clustering at that value of *r*. The maximum value of the *L(r) - r* function (*L_max_*) is an unbiased summary parameter that can be used to quantify the extent of clustering^40^. At least 12 PM sheets were imaged, analyzed, and pooled. To evaluate the differences between replicated points, a Bootstrap test was employed as described in Equation C:

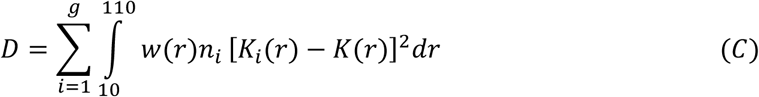

where *w(r)* = *r^-2^*, *K_i_(r)* = weighted mean K-function of the *i*^th^ group of size *n*_i_, and *K(r)* = combined weighted mean K-function of all the groups being compared (g). The observed value of D was ranked against 1000 Monte Carlo simulated values of D calculated using a set of residual K-functions derived from each *K_i_(r)*^40, 41^, the rank is an estimate of the probability that the null hypothesis of equivalence can be rejected.

### Peptide Synthesis

Peptides were synthesized using manual and automated solid-phase peptide synthesis **(Supplementary Figure 4)**. In a typical synthesis, Rink Amide resin SS 100–200 (0.300 g, 0.225 mmol; Advanced ChemTech) was swollen in DMF overnight, washed twice with DMF, and treated twice with 20% piperidine in DMF (2 × 10 min) to remove the initial Fmoc protecting group. Amino acid couplings were performed using 3 equivalents of Fmoc-protected amino acid, 2.85 equivalents of N,N,N′,N′-tetramethyl-O-(1H-benzotriazol-1-yl)uronium hexafluorophosphate (HBTU; Oakwood Products, Inc.), and 3 equivalents of N,N-diisopropylethylamine (DIEA) in 3 ml DMF. Coupling reactions proceeded at room temperature for 30 min, followed by resin filtration. A second coupling reaction (double coupling) was then performed, followed by capping with 200 μl acetic anhydride and 200 μL DIEA. The resin was subsequently treated twice with 20% piperidine in DMF (2 × 10 min), and additional amino acids were coupled as described above. Acetic anhydride capping was performed only after incorporation of the first amino acid. At the conclusion of synthesis, the resin was washed five times with DMF, five times with DCM, and dried under vacuum.

### Modeling

The peptide sequence was modeled using Pepfold4^42^, which generated a helical structure, consistent with the predicted structure of DIRAS3. FlexPepDock^43^ was used to dock this peptide to the structure of GMPPNP-loaded KRAS G12V (PDB entry 6GOE), starting from an approximate structure in which the peptide was proximal to KRAS switch I on the basis of the NMR chemical shift perturbations that we reported previously^19^. Both Cys residues were facing away from KRAS in the resulting model and were connected with the bis-phenyl linker using the PyMOL Molecular Graphics System (Version 3.1.6.1 Schrödinger, LLC.), which was also used to visualize the model.

### Confocal imaging

RASless MEFs expressing KRAS^G12D^ cells were seeded on sterile glass coverslips in 12-well plates and cultured overnight to reach ∼40–50% confluency. Cells were washed once with PBS and incubated with 25 µM fluorescein-labeled DIRAS3 peptide (DIRAS3 peptide-FL) or 25 µM fluorescein (FL) as a control, diluted in phenol red–free complete medium for 24 h at 37°C. Following incubation, cells were washed three times with PBS to remove unbound peptide/fluorophore, and counterstained with DAPI (1 µg/ml, 5 min) to visualize nuclei. Cells were fixed with 4% paraformaldehyde (PFA) in PBS for 15 min at room temperature, washed three times with PBS, and mounted in aqueous mounting medium. No detergent permeabilization was performed to preserve membrane-associated fluorescence. Images were acquired using a laser-scanning confocal microscope equipped with a 63× oil immersion objective. Fluorescein was excited at 488 nm, and DAPI at 405 nm. Detector gain and laser power settings were initially optimized using control samples and subsequently kept constant across all experimental conditions. Corresponding DIC images were acquired for each field to delineate cell morphology and plasma membrane boundaries.

### Reverse-phase protein array (RPPA) analysis

RASless MEF KRAS G12D and Q61L clones were treated with either control peptide or DIRAS3 peptide (15 μM) for 72 h. Cell lysis was performed on ice using lysis buffer containing T-PER reagent (Pierce, #78510), 450 mM NaCl, 1 mM sodium orthovanadate, Pefabloc (AEBSF), aprotinin, pepstatin A, and leupeptin. For lysis, 25 μl buffer was added per well of a 6-well plate at ∼70% confluency, or 15 μl per well of a 12-well plate. Cells were scraped, collected, and stored at −80°C until analysis. Samples were immobilized onto nitrocellulose-coated glass slides using a robotic microarray platform. Each sample was printed in technical triplicate (n=3) together with internal reference standards for quality assurance and quality control. Arrays were immunostained to quantify the expression and activation levels of 150 biochemically linked proteins and phosphoproteins involved in cancer-related signaling pathways, with a particular focus on PI3K and MAPK signaling networks. Signal intensities were analyzed using an in-house-developed pipeline, and a single normalized intensity value was generated for each sample following: (i) subtraction of nonspecific binding from secondary antibodies and staining reagents, (ii) normalization to total protein content determined by Sypro Ruby staining, and (iii) averaging across technical replicates. Internal reference standards were used to confirm that all samples fell within the assay’s linear dynamic range and to monitor assay performance.

### KRAS co-immunoprecipitation and immunoblotting

RASless MEFs expressing KRAS^G12V^ or KRAS^Q61L^ were treated with vehicle, RMC-7977 (10 nM), or DIRAS3 peptide (25 μM) for 4 h. Cells were washed with ice-cold PBS and lysed in cell lysis buffer (Cell Signaling Technology, #9803) supplemented with 10 mM MgCl₂ and freshly added protease and phosphatase inhibitors. Lysates were maintained on ice and clarified by centrifugation at 14,000 × *g* for 10 min at 4°C. Equal amounts of total protein (0.5 mg) were incubated with rabbit anti-KRAS antibody (Proteintech, 12063-1-AP; 2 μg) or an equivalent amount of normal rabbit IgG (Cell Signaling Technology, #2729) and Protein A magnetic beads (Cell Signaling Technology, #73778) at 4°C overnight. Beads were washed 3 times with ice-cold lysis buffer, and bound proteins were eluted in reducing SDS sample buffer. Immunoprecipitated proteins were resolved by SDS-PAGE and analyzed by immunoblotting for p110α, RAF1, and KRAS. Matched input lysates were analyzed for p-AKT, total AKT, p-ERK, total ERK, and vinculin. Immunoreactive bands were detected using HRP-conjugated secondary antibodies and chemiluminescence. The experiment was performed independently twice with similar results.

### Statistical analysis

All statistical analyses were performed using GraphPad Prism (GraphPad Software). Data are presented as mean ± SEM unless otherwise indicated. For comparisons between two groups, a two-tailed unpaired Student’s t-test was used. For experiments involving two independent variables, two-way analysis of variance (ANOVA) followed by appropriate post hoc multiple-comparison testing was performed, as indicated in the figure legends. All experiments were repeated at least three times (biological replicates), with each condition measured in technical triplicate. A P value < 0.05 was considered statistically significant.

## Acknowledgments

The authors would like to thank members of the Bast laboratory for their discussion and suggestions. The authors also thank the RAS Initiative at the Frederick National Laboratory for Cancer Research (FNLCR), an NCI-designated program operated by Leidos Biomedical Research, Inc. under contract with the National Cancer Institute, for providing RASless MEFs as validated reference reagents. This work was supported by National Cancer Institute R01 CA266187 (R.C.B and Z.L., PIs), R01 GM151280 (J.F.H. and J.L., PIs), the MD Anderson Ovarian SPORE P50 CA281701 (R.C.B. and A.K.S., PIs), the National Cancer Institute, Department of Health and Human Services, the Shared Resources of the MD Anderson CCSG grant NCI P30CA16672, the Cancer Prevention Research Institute of Texas RP140429 (R.C.B., PI), RP200166 (S.W.M. and Z.L., PIs), and RP250213 (J.F.H and J.L., PIs); and generous donations from the Ann and Henry Zarrow Foundation, the Roberson Endowment, and Stuart and Gaye Lynn Zarrow.

## Author Contributions

G.B. designed and performed experiments, analyzed data, and co-wrote the manuscript. H.Y. and W.M. carried out cell culture and validation experiments. R.B.G. performed nanoclustering experiments. V.K.B. and A.A.G. designed and performed MD dynamics experiments. S.Y.J. performed mass spectrometry analyses. M.P. and A.G. conducted RPPA analyses. J.F.H. provided key expertise in RAS nanoclustering. C.B.M. performed structural modeling and data interpretation. S.W.M. and M.N.U. contributed to peptide design, synthesis, structural modeling, and data interpretation. J.L. provided key expertise and critical guidance in RAS nanoclustering and contributed to the mechanistic interpretation of the results. F.M. provided critical feedback on the experimental findings, contributed valuable mechanistic insights and suggestions, and helped strengthen the overall presentation of the study. R.C.B. and Z.L. planned and supervised the study, provided resources and critical guidance, and contributed to the overall conceptual framework and manuscript preparation. All authors discussed the results, provided critical feedback, and approved the final manuscript.

## Data Availability

All data generated or analyzed during this study are included in this published article and its supplementary information files. Additional information supporting the findings of this study is available from the corresponding author upon request.

## References

1. Kim, D. et al. Pan-KRAS inhibitor disables oncogenic signalling and tumour growth. Nature 619, 160–166 (2023).

2. Hayes, T.K. et al. Long-Term ERK Inhibition in KRAS-Mutant Pancreatic Cancer Is Associated with MYC Degradation and Senescence-like Growth Suppression. Cancer Cell 29, 75–89 (2016).

3. Punekar, S.R., Velcheti, V., Neel, B.G. & Wong, K.K. The current state of the art and future trends in RAS-targeted cancer therapies. Nat Rev Clin Oncol 19, 637–655 (2022).

4. Sang, B. et al. Disrupted molecular glue complex drives RAS inhibitor resistance. Cell (2026).

5. Nissley, D.V., Stephen, A.G., Yi, M. & McCormick, F. Progress in Targeting KRAS Directly. Methods Mol Biol 2797, 1–12 (2024).

6. Simanshu, D.K., Philips, M.R. & Hancock, J.F. Consensus on the RAS dimerization hypothesis: Strong evidence for lipid-mediated clustering but not for G-domain-mediated interactions. Mol Cell 83, 1210–1215 (2023).

7. Venkatanarayan, A. et al. CRAF dimerization with ARAF regulates KRAS-driven tumor growth. Cell Rep 38, 110351 (2022).

8. Zhou, Y. & Hancock, J.F. RAS nanoclusters are cell surface transducers that convert extracellular stimuli to intracellular signalling. FEBS Lett 597, 892–908 (2023).

9. Luo, R.Z. et al. Genomic structure and promoter characterization of an imprinted tumor suppressor gene ARHI. Biochim Biophys Acta 1519, 216–222 (2001).

10. Sutton, M.N. et al. DIRAS3 (ARHI) Blocks RAS/MAPK Signaling by Binding Directly to RAS and Disrupting RAS Clusters. Cell Rep 29, 3448–3459 e3446 (2019).

11. Liang, X. et al. Membrane anchoring of the DIRAS3 N-terminal extension permits tumor suppressor function. iScience 26, 108151 (2023).

12. Bildik, G., Liang, X., Sutton, M.N., Bast, R.C., Jr. & Lu, Z. DIRAS3: An Imprinted Tumor Suppressor Gene that Regulates RAS and PI3K-driven Cancer Growth, Motility, Autophagy, and Tumor Dormancy. Mol Cancer Ther 21, 25–37 (2022).

13. Zhou, Y. et al. SIGNAL TRANSDUCTION. Membrane potential modulates plasma membrane phospholipid dynamics and K-Ras signaling. Science 349, 873–876 (2015).

14. Belanis, L., Plowman, S.J., Rotblat, B., Hancock, J.F. & Kloog, Y. Galectin-1 is a novel structural component and a major regulator of h-ras nanoclusters. Mol Biol Cell 19, 1404–1414 (2008).

15. Zhou, Y. et al. Lipid-Sorting Specificity Encoded in K-Ras Membrane Anchor Regulates Signal Output. Cell 168, 239–251 e216 (2017).

16. Klimpel, A., Stillger, K., Wiederstein, J.L., Kruger, M. & Neundorf, I. Cell-permeable CaaX-peptides affect K-Ras downstream signaling and promote cell death in cancer cells. FEBS J 288, 2911–2929 (2021).

17. Michaelson, D. et al. Postprenylation CAAX processing is required for proper localization of Ras but not Rho GTPases. Mol Biol Cell 16, 1606–1616 (2005).

18. Cox, A.D., Der, C.J. & Philips, M.R. Targeting RAS Membrane Association: Back to the Future for Anti-RAS Drug Discovery? Clin Cancer Res 21, 1819–1827 (2015).

19. Gray, J.P. et al. DIRAS3-derived cyclic peptides disrupt KRAS-RAF interaction and KRAS nanoclustering, suppressing KRAS-driven pancreatic and ovarian tumors. iScience 28, 113686 (2025).

20. Fleming, J. et al. AlphaFold Protein Structure Database and 3D-Beacons: New Data and Capabilities. J Mol Biol 437, 168967 (2025).

21. Ambrogio, C. et al. KRAS Dimerization Impacts MEK Inhibitor Sensitivity and Oncogenic Activity of Mutant KRAS. Cell 172, 857–868 e815 (2018).

22. Araya, M.K. et al. Differential Lipid Binding Specificities of RAP1A and RAP1B are Encoded by the Amino Acid Sequence of the Membrane Anchors. J Am Chem Soc 146, 19782–19791 (2024).

23. Remorino, A. et al. Gradients of Rac1 Nanoclusters Support Spatial Patterns of Rac1 Signaling. Cell Rep 21, 1922–1935 (2017).

24. Lechuga, C.G. et al. RASless MEFs as a Tool to Study RAS-Dependent and RAS-Independent Functions. Methods Mol Biol 2262, 335–346 (2021).

25. Tran, T.H. et al. KRAS interaction with RAF1 RAS-binding domain and cysteine-rich domain provides insights into RAS-mediated RAF activation. Nat Commun 12, 1176 (2021).

26. Baljuls, A. et al. The tumor suppressor DiRas3 forms a complex with H-Ras and C-RAF proteins and regulates localization, dimerization, and kinase activity of C-RAF. J Biol Chem 287, 23128–23140 (2012).

27. Holderfield, M. et al. Concurrent inhibition of oncogenic and wild-type RAS-GTP for cancer therapy. Nature 629, 919–926 (2024).

28. Marasco, M. et al. Direct Inhibition of RAS Reveals the Features of Oncogenic Signaling Driven by RAS G12 and Q61 Mutations. Cancer Discov 15, 1392–1409 (2025).

29. Navarro-Jimenez, M., Gonzalez, B., Mulet, N., Hierro, C. & Alonso, S. KRAS-targeted therapies in colorectal cancer: a systematic analysis of mutations, inhibitors, and clinical trials. NPJ Precis Oncol 9, 380 (2025).

30. Gebregiworgis, T. et al. The Q61H mutation decouples KRAS from upstream regulation and renders cancer cells resistant to SHP2 inhibitors. Nat Commun 12, 6274 (2021).

31. Wolpin, B.M. et al. Daraxonrasib in Previously Treated Advanced RAS-Mutated Pancreatic Cancer. N Engl J Med 394, 1790–1802 (2026).

32. Bildik, G. et al. DIRAS3 induces autophagy and enhances sensitivity to anti-autophagic therapy in KRAS-driven pancreatic and ovarian carcinomas. Autophagy 20, 675–691 (2024).

33. Rappsilber, J., Ishihama, Y. & Mann, M. Stop and go extraction tips for matrix-assisted laser desorption/ionization, nanoelectrospray, and LC/MS sample pretreatment in proteomics. Anal Chem 75, 663–670 (2003).

34. Abramson, J. et al. Addendum: Accurate structure prediction of biomolecular interactions with AlphaFold 3. Nature 636, E4 (2024).

35. Jo, S., Kim, T., Iyer, V.G. & Im, W. CHARMM-GUI: a web-based graphical user interface for CHARMM. J Comput Chem 29, 1859–1865 (2008).

36. Lee, J. et al. CHARMM-GUI Input Generator for NAMD, GROMACS, AMBER, OpenMM, and CHARMM/OpenMM Simulations Using the CHARMM36 Additive Force Field. J Chem Theory Comput 12, 405–413 (2016).

37. Hutchins, C.M. & Gorfe, A.A. Intrinsically Disordered Membrane Anchors of Rheb, RhoA, and DiRas3 Small GTPases: Molecular Dynamics, Membrane Organization, and Interactions. J Phys Chem B 128, 6518–6528 (2024).

38. Abraham, M.J., Murtola, T., Schulz, R., Páll, S., Smith, J. C., Hess, B., & Lindahl, E. GROMACS: High performance molecular simulations through multi-level parallelism from laptops to supercomputers. SoftwareX Volumes 1-2, Pages 19–25 (2015).

39. Shaw, D.E., Adams, P. J., Azaria, A., Bank, J. A., Batson, B., Bell, A.,…& Yuh, K. A. in SC ‘21: Proceedings of the International Conference for High Performance Computing, Networking, Storage and Analysis 1–11 (2021).

40. Plowman, S.J., Muncke, C., Parton, R.G. & Hancock, J.F. H-ras, K-ras, and inner plasma membrane raft proteins operate in nanoclusters with differential dependence on the actin cytoskeleton. Proc Natl Acad Sci U S A 102, 15500–15505 (2005).

41. Diggle, P.J., Mateu, J., & Clough, H. E. Comparison between parametric and non-parametric approaches to the analysis of replicated spatial point patterns. Advances in Applied Probability. Advances in Applied Probability 32(2), 331-343. (2000).

42. Rey, J., Murail, S., de Vries, S., Derreumaux, P. & Tuffery, P. PEP-FOLD4: a pH-dependent force field for peptide structure prediction in aqueous solution. Nucleic Acids Res 51, W432–W437 (2023).

43. Lyskov, S. et al. Serverification of molecular modeling applications: the Rosetta Online Server that Includes Everyone (ROSIE). PLoS One 8, e63906 (2013).

