## Supplementary data for "DIRAS3 broadly suppresses signaling from diverse oncogenic KRAS hotspot variants through an inhibitory interface"

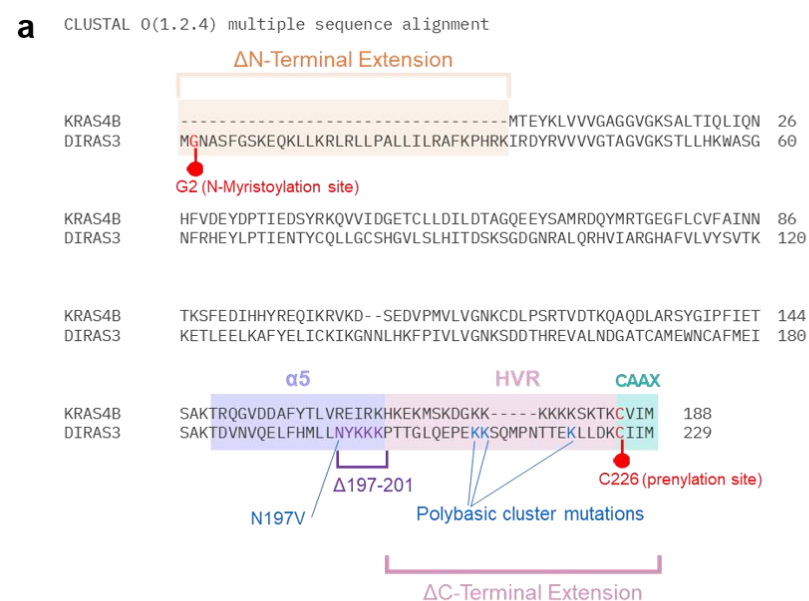

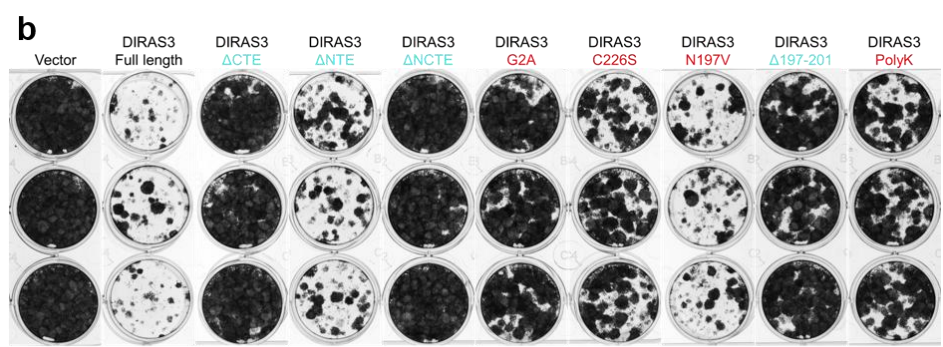

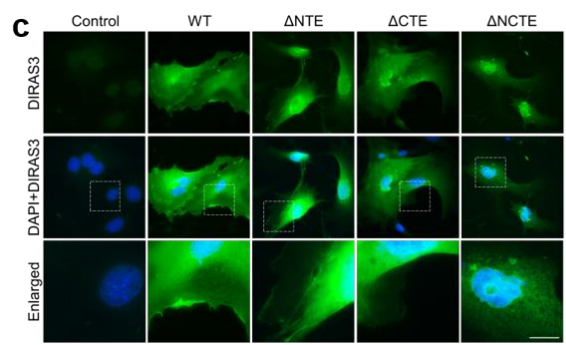

**Supplementary Fig. 1: Functional analysis of DIRAS3 deletion and point mutants.** Amino acid sequence alignment of DIRAS3 and KRAS4B, showing DIRAS3 deletion and point mutants targeting the N-terminal extension (ΔNTE), C-terminal extension (ΔCTE), combined termini (ΔNCTE), α5 helical region (Δ197–201 and N197V), and lipid-modifying residues (G2A and C226S), and the polybasic membrane-binding region (PolyK; K211Q/K212Q/K221Q). Mutated residues and deleted regions are indicated in the alignment. **b.** Representative triplicate images from the colony formation assay shown in Fig. 1D following transient expression of WT or mutant DIRAS3 in KRAS<sup>G12D</sup> RASless MEFs. Colonies were stained with Coomassie Brilliant Blue 10-14 d after seeding. **c.** Representative immunofluorescence images of WT DIRAS3 and NTD-, CTD-, and NCTD-deletion mutants expressed in KRAS<sup>G12D</sup> RASless MEFs. DIRAS3 is shown in green. Scale bar, 10 μm.

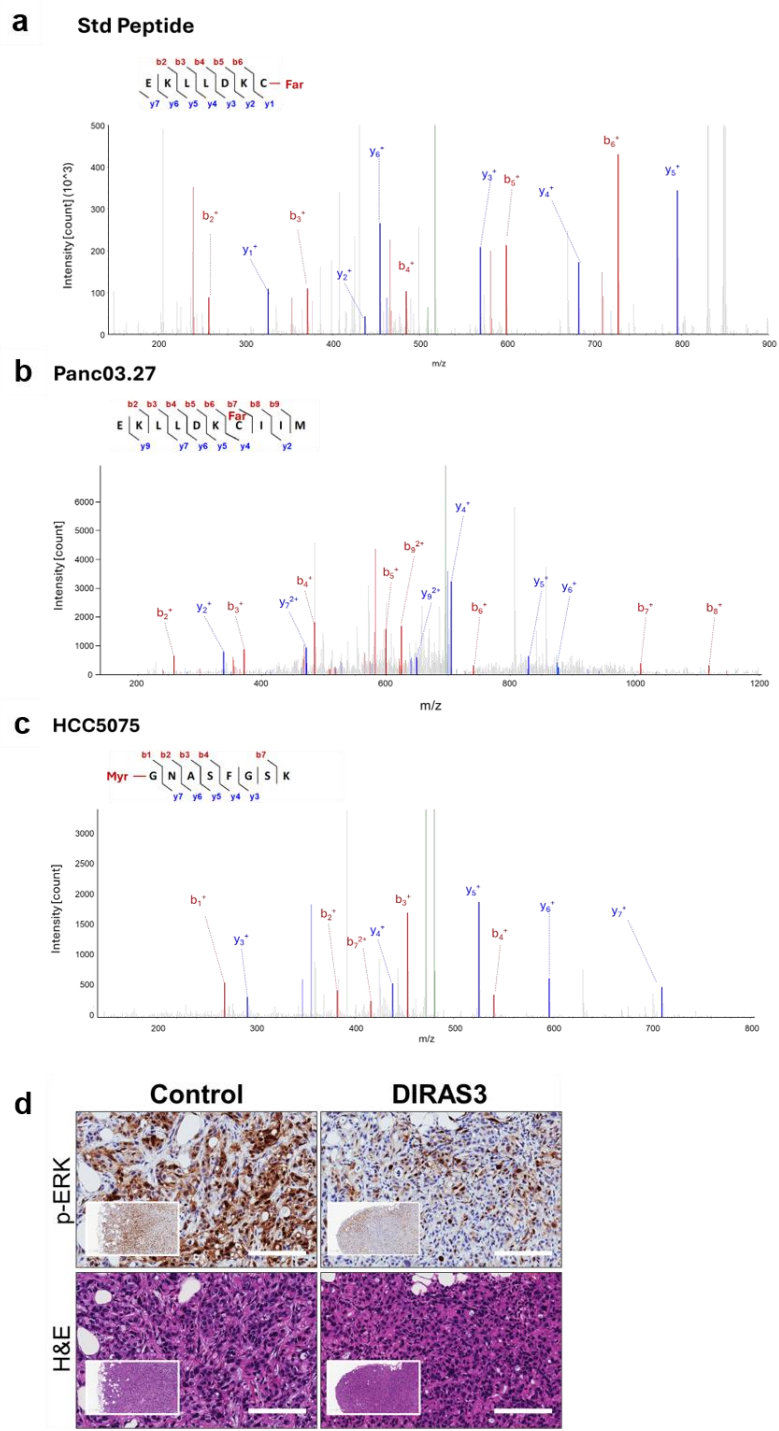

**Supplementary Fig. 2: Mass spectrometry identification of dual lipid modifications of DIRAS3 and suppression of p-ERK signal in vivo.** **a.** MS/MS spectrum of a standard farnesylated peptide used as a control. **b.** Identification of C-terminal lipid modification of DIRAS3. Mass spectrometry detected the peptide EKLLDKCIIM containing residue C226 from Panc03.27 pancreatic ductal adenocarcinoma cells with dox-induced DIRAS3 expression. DIRAS3 was enriched using an anti-DIRAS3 polyclonal antibody, followed by reduction, alkylation, trypsin digestion, and LC-MS/MS analysis. **c.** Identification of N-terminal lipid modification of DIRAS3. Mass spectrometry detected the peptide GNASFSGK containing residue G2 from HCC5075 low-grade serous ovarian cancer cells with dox-induced DIRAS3 expression. DIRAS3 was enriched using an anti-DIRAS3 polyclonal antibody, followed by reduction, alkylation, trypsin digestion, and LC-MS/MS analysis. **d.** Representative hematoxylin and eosin (H&E) and immunohistochemical staining of p-ERK in tumor sections from AsPC-1 xenografts with control or doxycycline-induced DIRAS3 re-expression. Representative images are shown. Scale bar: 40 $\mu\text{m}$.

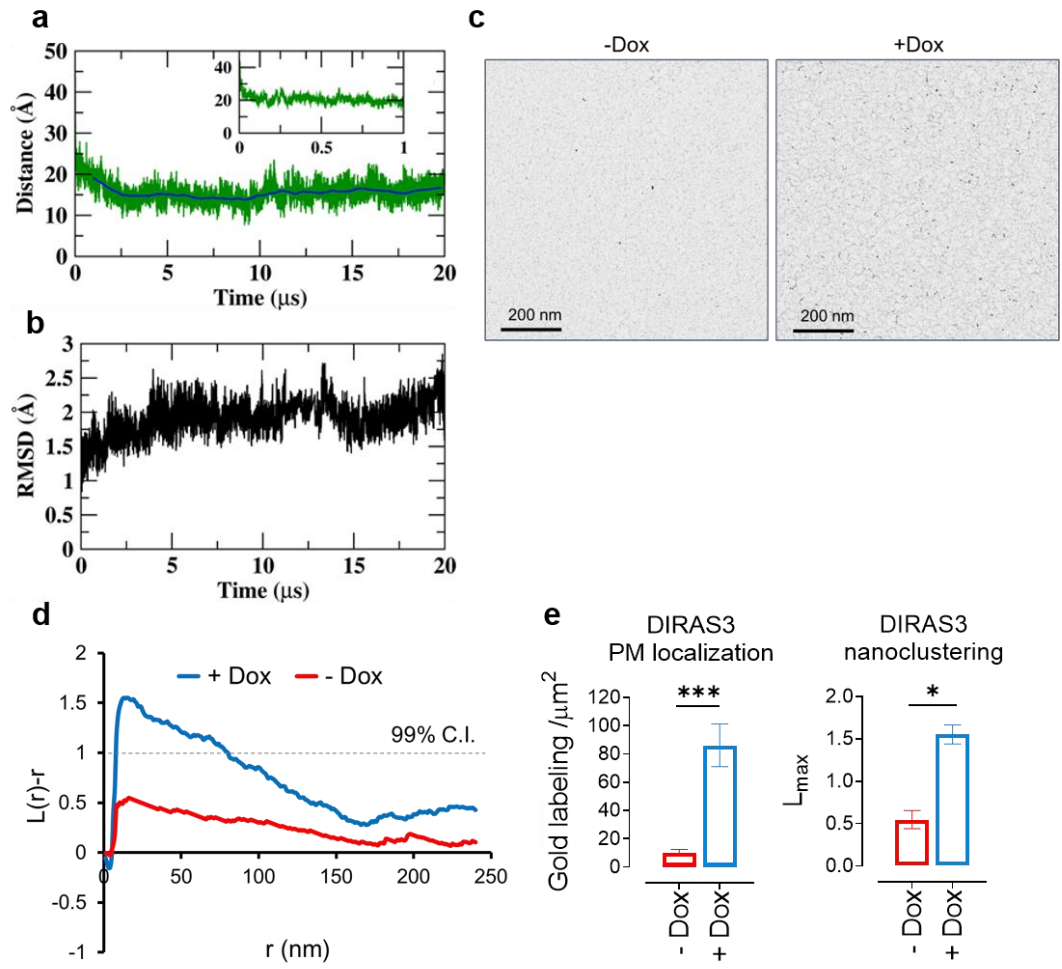

**Supplementary Fig. 3: Molecular dynamics simulations and electron microscopy analyses of DIRAS3 membrane association and plasma membrane nanoclustering.** **a.** Distance between the G-domain and phosphate midplane of the inner leaflet over time. The blue line represents the running average, and the inset shows rapid membrane recruitment during the first 1 $\mu\text{s}$ of the simulation. **b.** Root Mean Square Deviation (RMSD) of the Cα atoms of the G-domain relative to the starting structure, indicating rapid equilibration and conformational stability. **c.** Representative images (1 $\mu\text{m}^2$) of 6 nm-gold-anti-DIRAS3 labelled PM sheets of HCC5075 cells with (+Dox) or without (-Dox) doxycycline-induced expression of DIRAS3 were imaged with the EM at 100,000X magnification. **d.** Plots of standardized linear K-functions were shown for control and DIRAS3-expressing HCC5075 cells. **e.** Quantification of plasma membrane localization as gold-labeling density and nanoclustering as the maximum value of $L(r)-r$ ($L_{\text{max}}$). Bootstrap testing was used to evaluate statistical differences in the extent of nanoclustering (\*$p < 0.05$). Gold-labeling density was compared using a two-tailed Student's t-test (\*\*$P < 0.001$), and $L_{\text{max}}$ was compared by bootstrap analysis ($P < 0.05$).

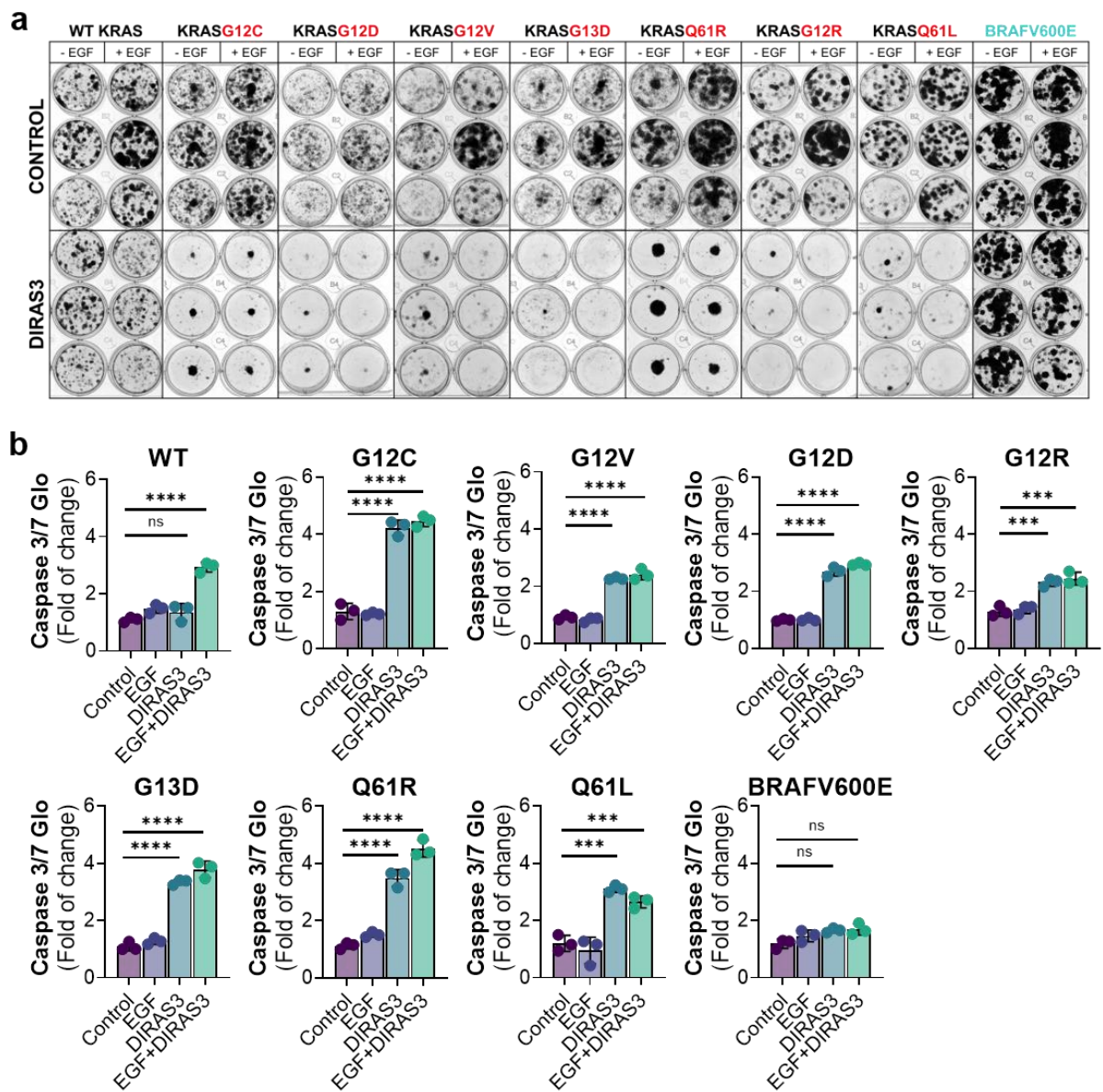

**Supplementary Fig. 4: Effects of DIRAS3 expression on colony formation and apoptosis in KRAS-mutant RASless MEFs.** **a.** Representative triplicate images of colony formation assays shown in Fig. 2E following transient expression of full-length DIRAS3 in KRAS-mutant RASless MEFs cultured in the presence or absence of EGF (20 ng/mL). Colonies were stained with Coomassie Brilliant Blue 10-14 d after seeding. **b.** Caspase-3/7 Glo assay measuring apoptosis in KRAS-mutant MEFs following DIRAS3 expression.

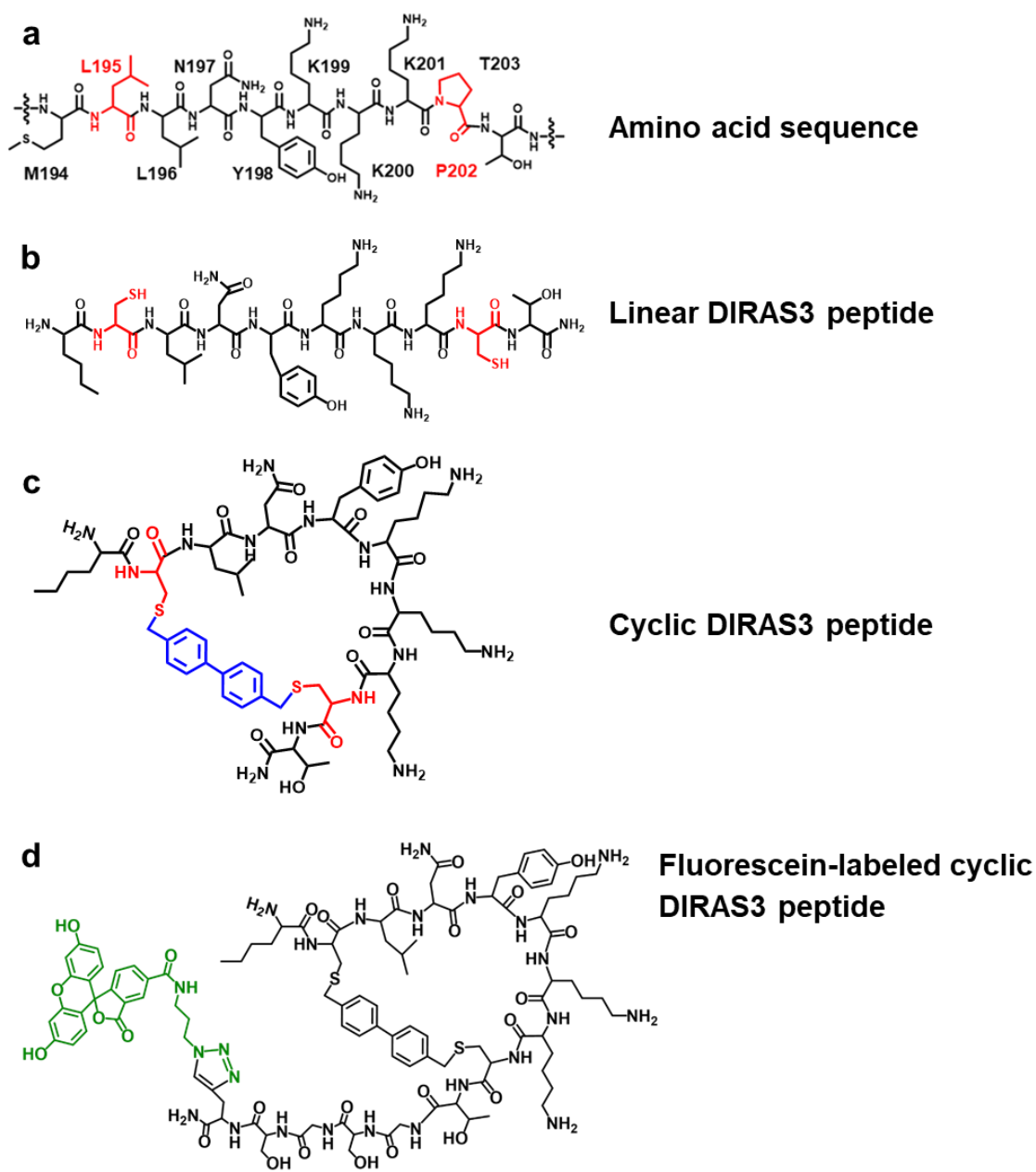

**Supplementary Fig. 5: Sequence and structural design of DIRAS3-derived peptides.** **a.** Amino acid sequence corresponding to residues 194-203 of DIRAS3. **b.** Linear DIRAS3 peptide derived from the $\alpha 5$ helical region. **c.** Bisphenyl-cyclized DIRAS3 $\alpha 5$ peptide. **d.** Fluorescein-labeled version of bisphenyl-cyclized DIRAS3 peptide.

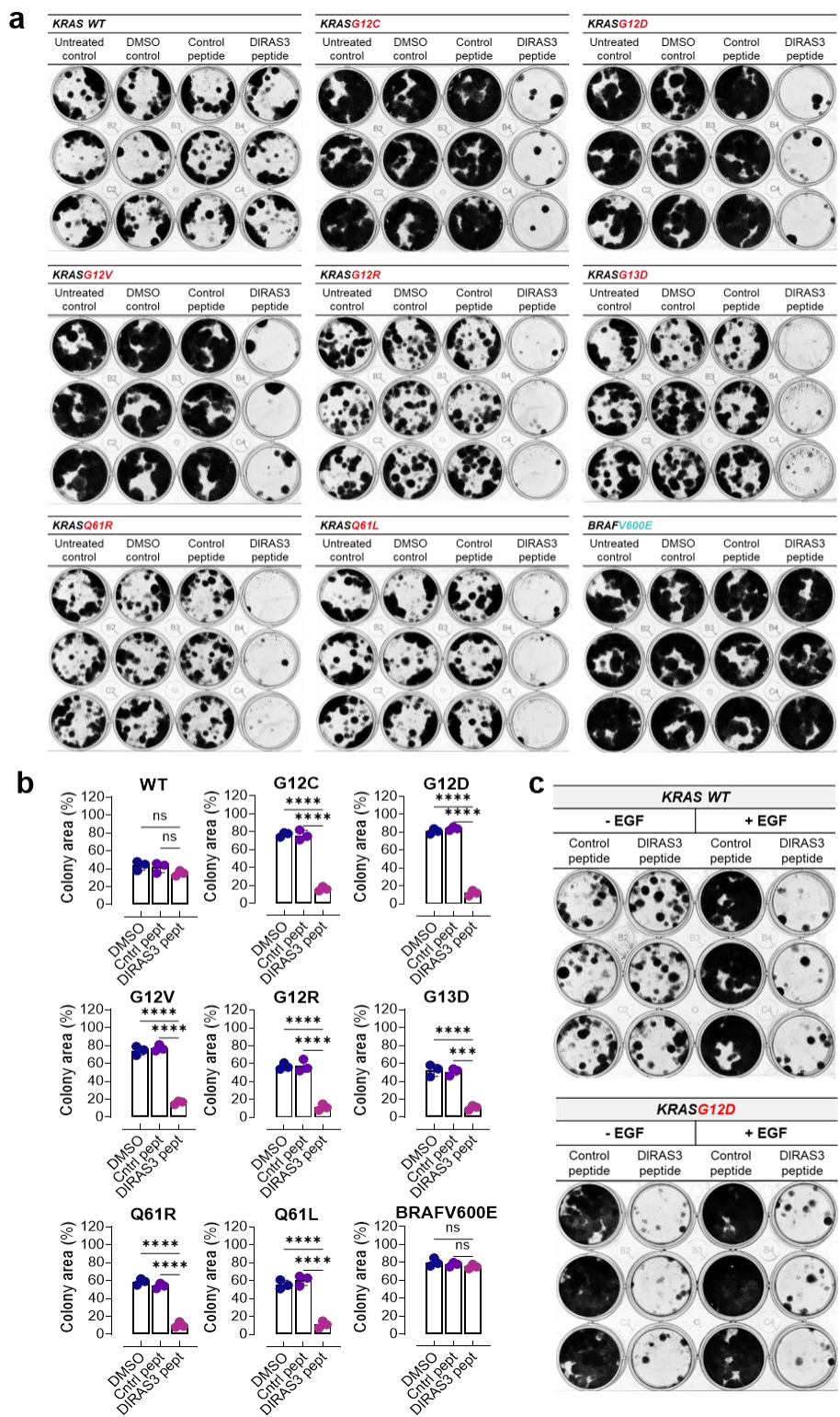

**Supplementary Fig. 6. DIRAS3 peptide suppresses clonogenic growth in RASless MEFs.** **a.** Representative triplicate images of colony formation assays shown in Fig. 3D, performed in KRAS-mutant RASless MEFs treated with DIRAS3 peptide (10 μM) or control peptide (10 μM). Colonies were stained with Coomassie Brilliant Blue 10–14 d after seeding. **b.** Quantification of colony area relative to the control peptide shown in panel a. Data are presented as mean ± SD from three independent biological experiments ($n = 3$). Statistical significance was determined by two-way ANOVA. ns, $p > 0.05$; \*\*\*$p < 0.001$; \*\*\*\*$p < 0.0001$. **c.** Representative triplicate images of colony formation assays following treatment with DIRAS3 peptide (10 μM) in KRAS WT and KRAS<sup>G12D</sup> RASless MEFs cultured in the presence or absence of EGF (20 ng/mL). Colonies were stained with Coomassie Brilliant Blue 10–14 d after seeding.

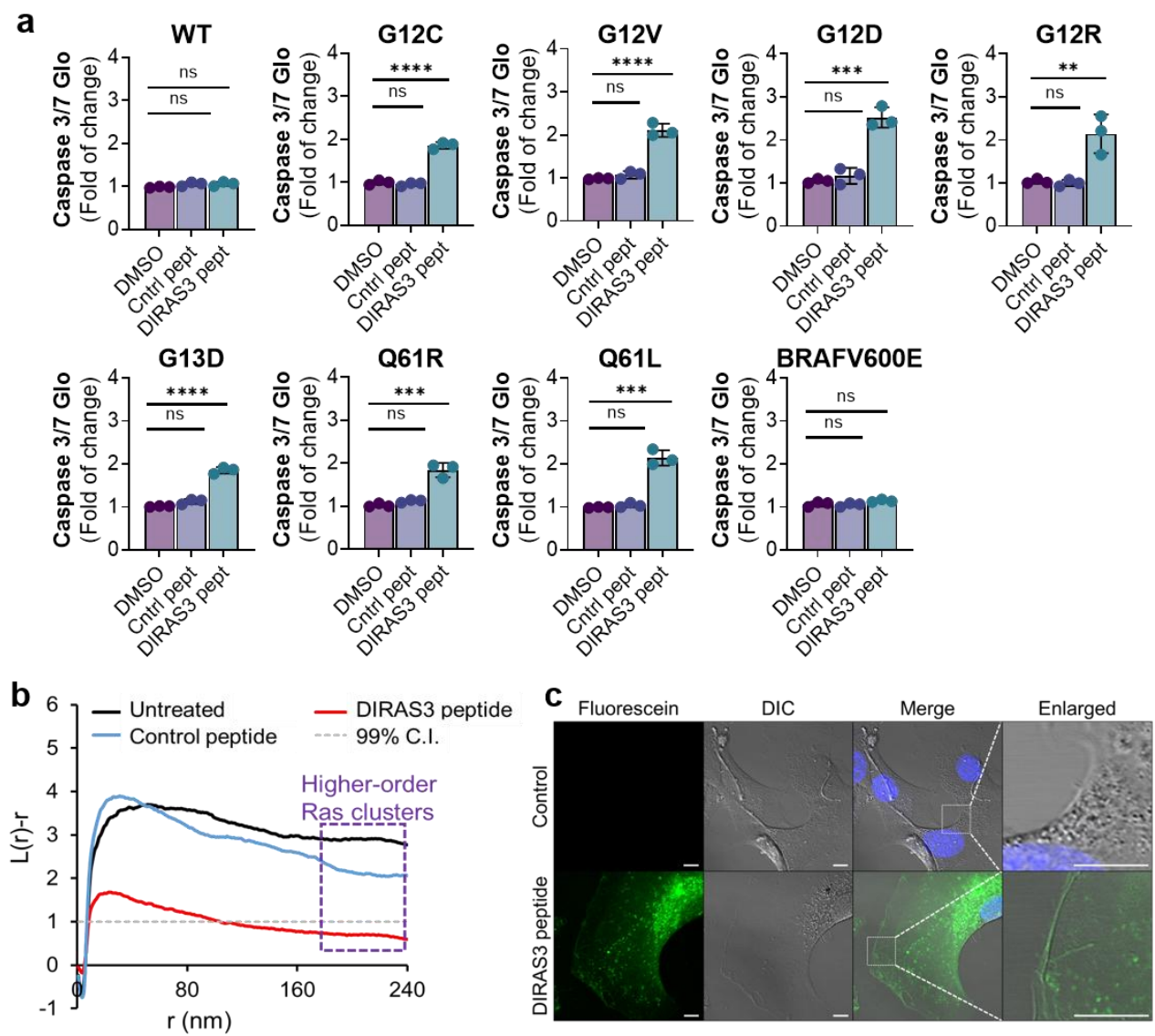

**Supplementary Fig. 7: Activity and cellular localization of the DIRAS3 $\alpha 5$ mimetic peptide.** **a.** Caspase-3/7-Glo assay measuring apoptosis in KRAS-mutant RASless MEFs following treatment with cyclic DIRAS3 peptide (15 μM). Data are presented as mean ± SD from three independent biological experiments ($n = 3$). **b.** Representative standardized linear K-function plots of GFP-KRAS<sup>G12V</sup> nanoclustering following treatment with DIRAS3 peptide, control peptide, or no treatment. Corresponding quantification is shown in Fig. 3E. **c.** Representative confocal images of KRAS<sup>G12D</sup> RASless MEFs treated with fluorescein-labeled cyclic DIRAS3 peptide (DIRAS3-FL) or control fluorophore. Nuclei were counterstained with DAPI. Differential interference contrast (DIC) images show cell morphology. Representative of three independent experiments. Scale bar, 10 μm.

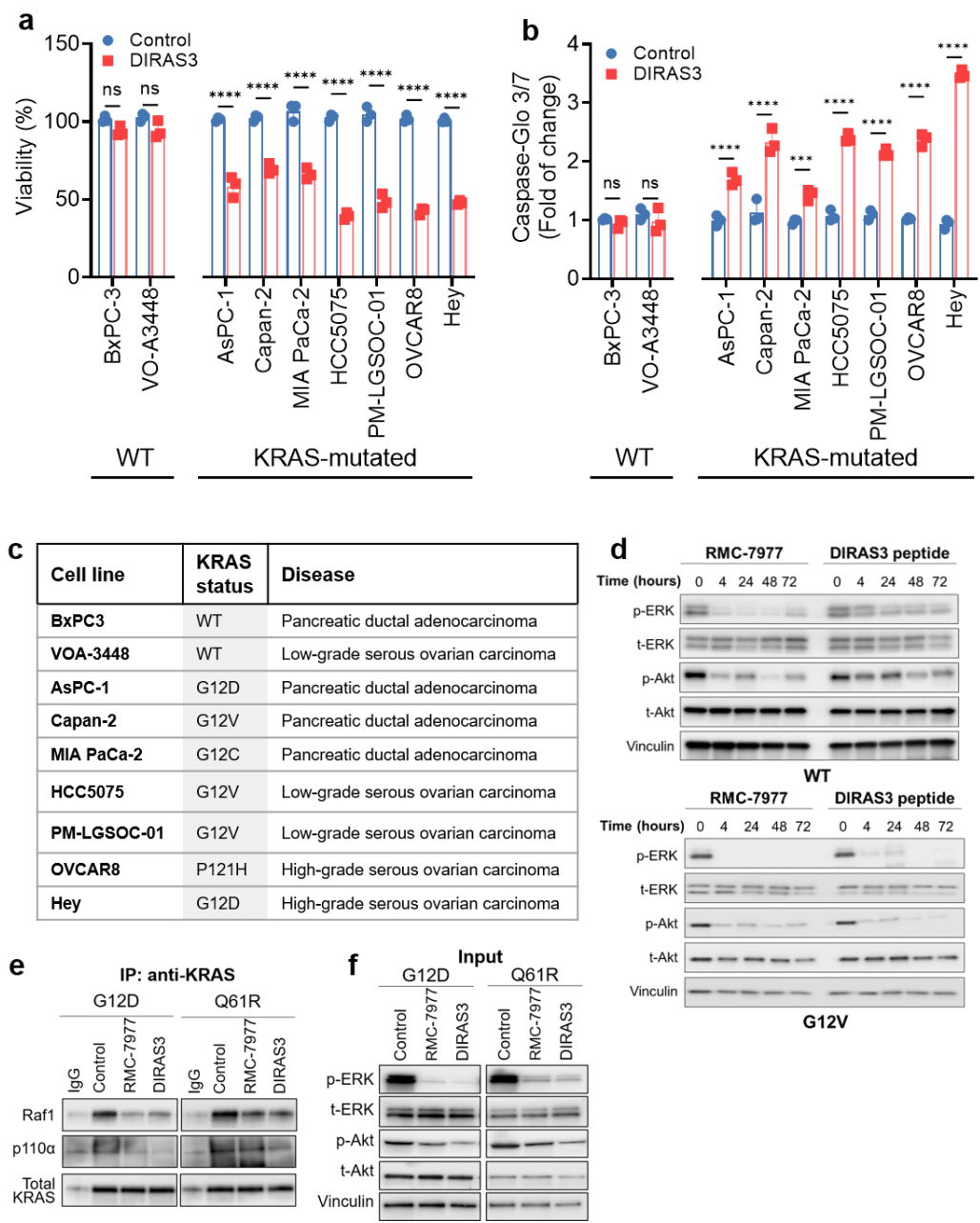

**Supplementary Fig. 8: Activity of the DIRAS3 $\alpha 5$ mimetic peptide across KRAS-mutant cancer cell models.** **a.** Cell viability of pancreatic and ovarian cancer cell lines harboring WT KRAS or oncogenic KRAS hotspot variants following treatment with control peptide or DIRAS3 $\alpha 5$ mimetic peptide (15 μM) for 96 h. Cell viability was measured by CellTiter-Glo. **b.** Caspase-3/7-Glo assay measuring apoptosis in cancer cell lines treated with control peptide or DIRAS3 $\alpha 5$ mimetic peptide (15 μM) for 96 h. Data are presented as mean ± SD from three independent biological experiments ($n = 3$). Statistical testing was performed by two-way ANOVA. The columns indicate the mean, and the bars indicate the SD (ns: $p > 0.05$, \*\*$p < 0.01$, \*\*\*$p < 0.001$, \*\*\*\*$p < 0.0001$). **c.** Characteristics of the cancer cell line panel used in this study, including tissue origin and KRAS mutation status. **d.** Representative immunoblot analysis of MAPK (phospho-ERK1/2) and PI3K/AKT (phospho-AKT) signaling in RASless MEFs rescued with KRAS WT and KRAS<sup>G12D</sup> following treatment with RMC-7977 or the DIRAS3 peptide. Representative blots from three independent biological experiments are shown. **e.** KRAS-effector interactions in RASless MEFs expressing KRASG12D or KRASQ61R following treatment with vehicle, RMC-7977 (10 nM), or DIRAS3 peptide (25 μM) for 4 hours. KRAS was immunoprecipitated from 500 μg of whole-cell lysates, and associated p110α and RAF1 were detected by immunoblotting; immunoprecipitated KRAS served as the loading control. Blots are representative of 2 independent biological replicates. **f.** Immunoblots of the corresponding input lysates from the experiments in (e), probed for phospho-ERK1/2, total ERK1/2, phospho-AKT, total AKT, and Vinculin. Loss of RAS–RAF1 association in (e), together with reduced ERK phosphorylation here, confirms target engagement by RMC-7977 at the concentration used. Blots are representative of 2 independent biological replicates.
